# PSD-95 drives binocular vision maturation critical for predation

**DOI:** 10.1101/2025.10.22.683936

**Authors:** Subhodeep Bhattacharya, Livia J.F. Wilod Versprille, Jonathan Ho, Xiaojie Huang, Cornelia Schöne, Oliver M. Schlüter, Siegrid Löwel

## Abstract

Postsynaptic density protein 95 (PSD-95) is a signalling scaffold within the postsynaptic density of excitatory synapses which drives silent synapse maturation during critical periods (CP). Binocularity develops during visual CPs and matures before its closure. Despite lifelong critical period plasticity, PSD-95 knock-out (KO) mice were reported with only subtle sensory phenotypes as adults. To assess PSD-95’s role in ethologically relevant binocular visual processing, we compared prey capture behaviour in PSD-95 KO and wild-type (WT) mice. KO mice were profoundly impaired in diverse epochs of predatory behaviour, but exhibited improved prey localisation under monocular conditions, reminiscent of impaired binocular integration. This was confirmed in an orientation discrimination task, where KO mice were impaired binocularly but performed monocularly like WT mice. Furthermore, binocular orientation discrimination was impaired after knocking down PSD-95 in primary visual cortex (V1), while a superior colliculus-specific knock-down had no effect. These results support an essential role of PSD-95 in binocular behaviour, which becomes evident under ethologically demanding behaviours.

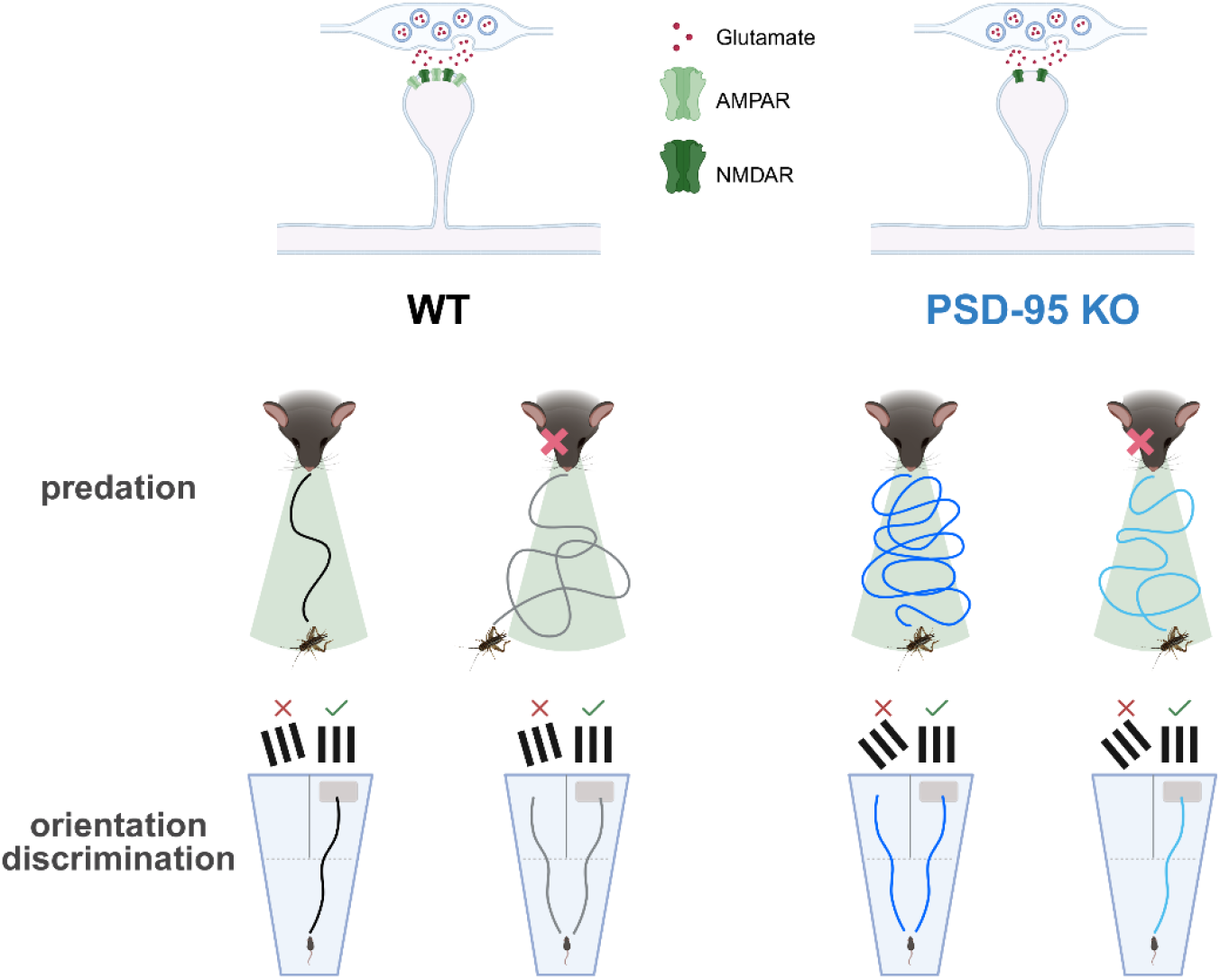

## INTRODUCTION

In their natural ecological context, mice rely on vision for a broad range of behaviours that are often not represented in standard laboratory settings^1^. Despite their relatively poor visual acuity, mice use vision for numerous natural activities such as avoiding predators, capturing prey, and navigating their environment. They are opportunistic feeders and hunt insects for nutritional enrichment, which as invaders can cause extinction of endemic populations^2–5^. Some mouse species, like the grasshopper mouse are specialized insect hunters, indicating predation as a highly ethologically relevant behaviour which substantially utilises the visual system for prey localisation^6^. Binocular vision is considered to support predatory hunting across species, including for non-foveate animals like mice, whose eyes are laterally placed^7,8^. Binocular vision provides depth cues for optimal prehensile movements in humans^9^ and can potentially break camouflage, improve predator-prey distance estimation and increase sensitivity in low contrast and low light conditions^10,11^. Previous studies assessing prey capture behaviour have observed that lab-bred adult wild-type (WT) mice naïve to crickets i) show a natural tendency to catch and consume them from the first day of exposure^3,12,13^, ii) require binocular vision for optimal hunting performance^14–16^ and iii) get better with more experience of prey-related salient visual features^17^. Juvenile WT mice during early visual critical periods (CP) hunt less efficiently than mature mice^18,19^, characterised by their inability to keep their head direction within the binocular visual field during active approaches towards prey^18^. In contrast, adult WT mice exhibit a strong binocular visual field bias during approaches^13,18^. It remains unresolved whether this bias, observed only in adult mice, is a result of critical period refinements in the visual cortex.

Neurons in the mouse primary visual cortex (V1) encode different properties of the visual scene including the ability to distinguish different orientations and spatial frequencies of a visual stimulus^20,21^ and are deemed to encode binocular visual information^22,23^. Notably, blocking V1 activity led to deficits in prey capture behaviour in adult mice^24^. Inefficient predatory hunting during visual CP correlates with a greater degree of binocular mismatch at the neuronal level in V1 before and during visual CP, which progressively improves with visual experience through CP and are well matched by early adulthood^25,26^. However, as these observations are correlative, it remains elusive whether these or other proper neural network refinements during CP sets up binocular properties for efficient prey hunting. During CP, V1 undergoes profound visual experience-dependent changes wherein relevant neuronal responses to visual stimuli are optimised to establish specialized visual abilities such as binocularity^22,23,25,26^. The connectivity between principal neurons is refined^27,28^, characterised by new synapse formation, their maturation, and elimination of unfavourable connections^29,30^. Silent synapses are nascent synapses lacking α-amino-3-hydroxy-5-methyl-4-isoxazolepropionic acid receptors (AMPAR) and serve as plasticity substrates during critical periods^28,31–33^. The postsynaptic signalling scaffold PSD-95 drives silent synapse maturation^34,35^ but PSD-95 overexpression hardly has any effect in mature neurons, when silent synapses are relatively sparse^36^, indicating a predominant role of PSD-95’s function during CPs. Without PSD-95, silent synapse maturation is impaired, and silent synapses are reinstated with knock-down of PSD-95 in the mature V1^32,37^. Consequently, the closure of the CP for ocular dominance plasticity (ODP), a classical procedure to test CP plasticity, is impaired in the absence of PSD-95 in V1 during postnatal development, reinstated when PSD-95 is knocked down in mature V1, and leads to lifelong juvenile-like ODP in PSD-95 knock-out (KO) mice^32^. In addition, loss of PSD-95 causes structural changes in the adult mouse V1 which are reminiscent of those observed in juvenile WT mice, i.e., enhanced spine elimination on the apical dendrites of layers 2/3 pyramidal neurons following monocular deprivation^38^. These and other results established PSD-95-dependent silent synapse maturation as key substrates for CP plasticity.

Both knock-out and knock-down of PSD-95 has no influence on V1 retinotopy and on general visual processing such as the optomotor reflex^32^. Indeed, the functional consequences on visual perception with loss of PSD-95 were comparatively subtle^34^. The visual water task (VWT), a form of forced-choice visual discrimination task, has been utilised to assess visual acuity^39^ and orientation discrimination^34^. VWT-based visual acuity was compromised in mice after monocular deprivation (MD) during CP^40^ and bilateral lesions in V1^41^. In VWT, compared to WT, PSD-95 KO mice displayed normal visual acuity but reduced orientation discrimination capabilities^34^. However, such a measure of orientation discrimination ability lacks behavioural relevance and whether a deficiency of PSD-95 translates into poorer behavioural outcomes in ethological contexts where binocular visual perception plays a decisive role, is unknown. Therefore, we assessed the role of vision, particularly binocular vision, by comparing PSD-95 KO and WT mice using the ethologically relevant prey capture task^13,14,18,19^. To probe the differential contribution of V1 and superior colliculus (SC), we assessed orientation discrimination in the VWT following local PSD-95 knock-down (KD). We hypothesised that impaired silent synapse maturation caused by loss of PSD-95 in V1 would compromise binocular vision and consequently, predatory behaviour.

We therefore tested binocular and monocular hunting performance in adult PSD-95 KO and WT mice. Tested binocularly, PSD-95 KO mice exhibited pronounced deficits in prey capture compared to WT mice. In contrast, compared to the binocular condition, KO mice displayed *enhanced* monocular prey capture performance, suggesting disturbed binocular processing. A similar phenotype was observed in the visual water task: KO mice had decreased binocular orientation discrimination compared to WT mice, but, in contrast to WT mice, improved during monocular vision to match performance of monocular WT mice. Notably, binocular orientation discrimination was selectively impaired only after a V1-specific, but not after a superior colliculus-specific PSD-95 knock-down. These findings demonstrate that the development and refinement of binocular visual ability, in terms of orientation discrimination and predatory behaviour, is strongly dependent on PSD-95-mediated maturation of AMPAR-silent synapses.

## RESULTS

### PSD-95 KO mice take longer to catch prey yet improve with monocular vision

While hunting behaviour in mice is instinctive, adult mice, compared to juveniles, are more efficient during initial hunting attempts^13,18^, suggesting that different motor and sensory features require experience-dependent refinement for optimal predatory behaviour. We tested the requirement of PSD-95-dependent CP binocular vision refinement for efficient predation by observing predatory behaviour of both PSD-95 WT and KO mice using an overhead camera in a behavioural arena^13^ (*Figure 1A*). Mice underwent three days of acclimatisation (A1-A3) and then caught crickets in the arena until consistent performance was achieved (D1-D_n-1_), followed by two test days under binocular (D_n_) and monocular (D_n+1_) conditions (*Figure 1B*). On the first training day, while 7/8 WT mice caught crickets at least in one of the trials, none of the PSD-95 KO mice caught crickets. KO animals took considerably more days than WT mice to successfully and reliably catch crickets (*Figure 1D*). On the first hunting day in the arena, KO mice displayed profound deficits in various epochs of hunting behaviour characterised by reduced approach rates, longer prey detection times, a diminished probability to maintain close proximity to prey during active approaches, greater proportion of inactive phases and decreased speed modulation (*Figure S1*). At the stage where mice exhibited consistent capture performance (D_n-1_), KO mice needed ∼5x longer to catch crickets than WT mice (*Figure 1E*). Comparison of capture times between the last training day (D_n-1_) and the binocular test day (D_n_) demonstrated that capture times were stable prior to the test phase in both WT and KO mice (*Figure 1F-G*).

**Figure 1.**
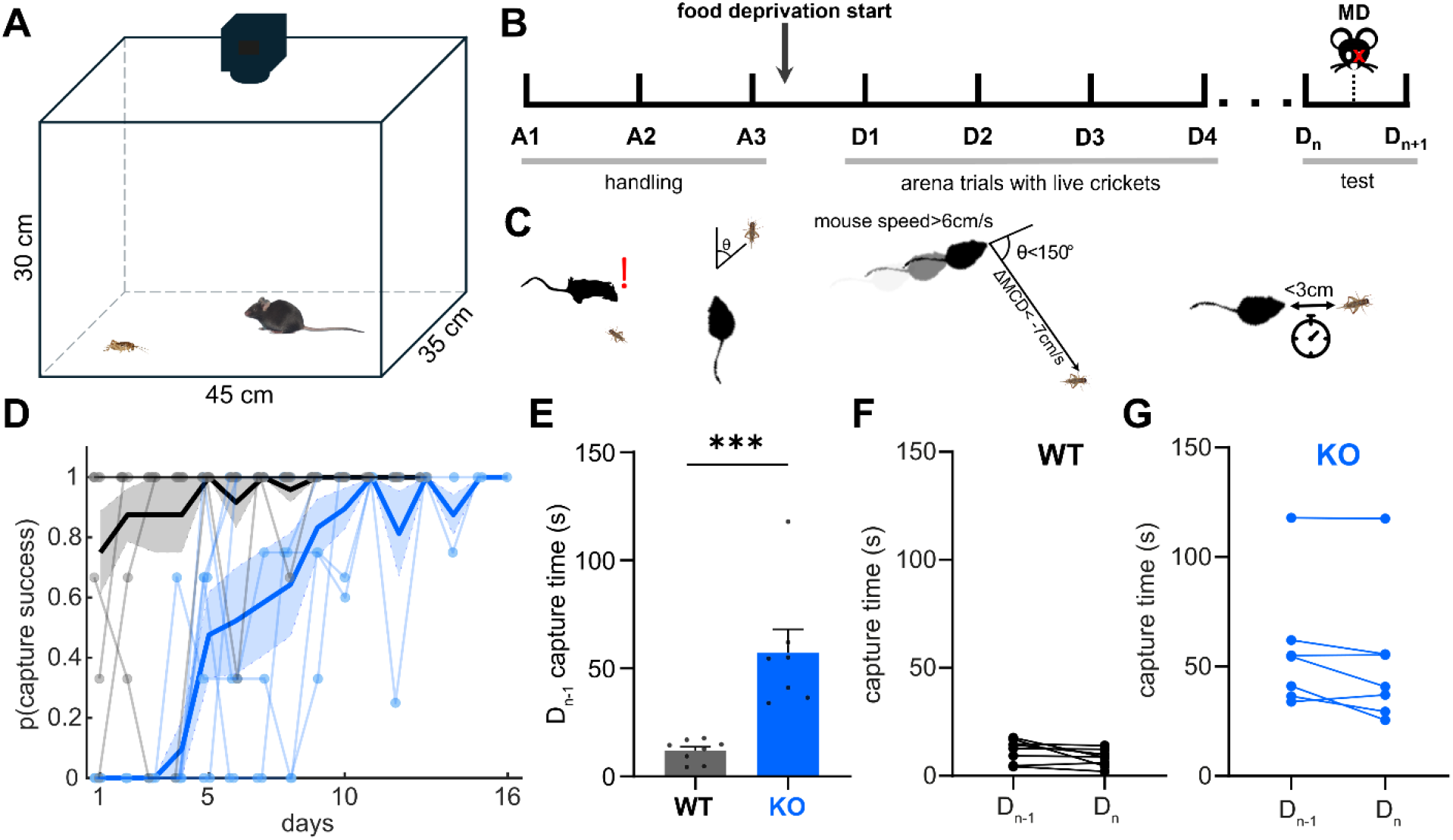
PSD-95 KO mice exhibit deficits in prey capture behaviour. (A) Schematic of the arena used for testing prey capture behaviour. Behaviour was recorded at 60 fps using an overhead camera (for details see Methods). (B) Experimental timeline with relevant phases indicated by grey lines; after 3 days of handling/acclimatisation (A1-A3), mice were food deprived and made to hunt crickets in the arena until they exhibited stable capture performance (D1-Dn-1). The following two test days consisted of mice hunting crickets first with intact binocular vision (Dn), after which one eye was sutured shut to allow monocular testing in the same animals ∼20-24 hrs later (Dn+1). (C) Schematic representation of predatory behaviour and parameters recorded. The first schematic depicts the time to 1^st^ approach, i.e. the time it takes for the mouse to detect the cricket and make its first approach. The second schematic depicts the azimuth between mouse and cricket, i.e. the head position of the mouse in relation to the cricket. The third schematic depicts an approach, defined as an epoch where the mouse speed was >6 cm/s, mouse-cricket distance decreased by a rate of 7 cm/s and the azimuth was <150 degrees. The fourth schematic depicts a mouse-cricket contact, defined as a mouse-cricket distance <3 cm and azimuth <90°. (D) Probability to catch crickets on a given day within 10 minutes of prey exposure. Individual data points are overlaid on the mean±SEM lines for each genotype. (E) Prey capture times of PSD-95 WT and KO mice on the last training day (Dn-1). Each point represents the median capture time of individual mice whereas the bars indicate the mean±SEM. (WT: 11.8±1.8 s, n=8; KO: 57.2±10.9 s, n=7; Mann-Whitney, p= 0.0003). (F and G) Comparison of median prey capture times for individual PSD-95 WT and KO mice between the last training day and the binocular test day, respectively (WT: Dn – Dn-1 = −3.6±1.71 s, n=8; Wilcoxon’s test, p= 0.055; KO: Dn – Dn-1 = −5.6±2.7 s, n=7; Wilcoxon’s test, p= 0.125).

Mice reliably capture prey by utilising binocular vision^14^ with the binocular representation of visual features, at the level of individual V1 neurons, undergoing development and refinement during CP^22,23,25,26^. The CP for ODP does not close in PSD-95 KO mice^32^, therefore we hypothesised that the observed deficit in binocular prey capture behaviour in the KOs might be due to specific disruptions in binocular visual abilities. To test to what extent binocular vision is required for prey capture behaviour, mice underwent monocular eye closure to compare binocular with monocular prey capture behaviour in the same animals (*Figure 2A-C*). Using monocular vision, individual capture times of WT mice increased by over 150%, whereas capture times of KO mice decreased by ∼12% (*Figure 2B*). By pooling and comparing individual trials between genotype and eye conditions using a linear mixed-effects model to account for inter-subject differences and trial-to-trial variability (LME; see *Table 1 and S1*), it was revealed that KO mice with binocular vision took ∼9x longer than WT mice to catch crickets (*Figure 2C*). Consistent with previous reports^14,16^, WT mice took >2x longer to successfully catch crickets under monocular conditions, thereby underscoring the importance of binocular vision for prey capture. In contrast, in PSD-95 KO mice, monocular vision resulted in a ∼2-fold *improvement* in capture times, indicating that binocular vision was compromised in PSD-95 KO mice and that primarily impaired vision, rather than other sensory modalities, contributed to the hunting impairment in PSD-95 KO mice.

**Figure 2.**
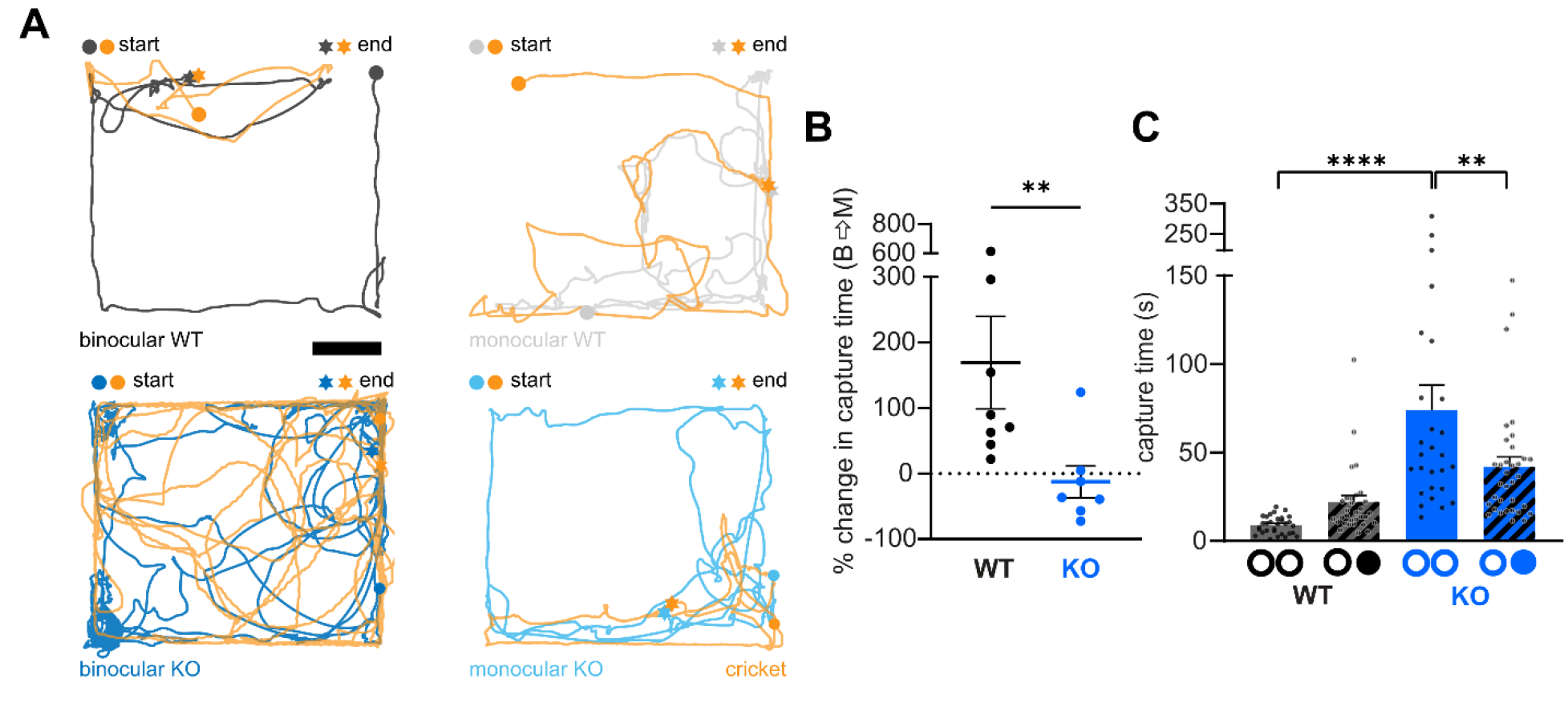
Monocular PSD-95 KO mice hunt faster than with intact binocular vision while WT mice get worse with monocular vision. (A) Example tracks of the same PSD-95 WT and KO mice under binocular and monocular condition, respectively, until successful capture (scale bar, 10 cm). (B) Percentage change in median capture times of individual WT/KO mice between binocular and monocular condition (WT: 169.2±70.5 %, n=8; KO: −12.7±24.8 %, n=7; Mann-Whitney test, p=0.006). (C) Within-group comparisons of prey capture times of WT and KO mice under both binocular and monocular (hatched bars) conditions, respectively. Each datapoint represents a trial and each bar represents the mean±SEM (WT binocular: 8.9±0.9 s, n=31; WT monocular: 22.2±3.6 s, n=32; KO binocular: 74.2±13.9 s, n=27; KO monocular: 42±5.6 s, n=34). ○○, binocular ○●, monocular. Linear mixed effects model with estimated marginal means for multiple comparisons (see Table 1 and S1). **p< 0.01 and ****p < 0.0001.

**Table 1.** Statistics of prey capture parameters in the light trials. Summary of the statistical outcomes of the type II Wald χ2 tests for each prey capture parameter. A complete overview of the statistical results can be found in Supplementary Table S1. Abbreviations: n.s. = non-significant; tr. = travelled.

| Parameter | Genotype |  | Eye condition |  | Genotype x eye condition |  |
| --- | --- | --- | --- | --- | --- | --- |
| | $\chi^2$(df) | p-value | $\chi^2$(df) | p-value | $\chi^2$(df) | p-value |
| Capture time | 19.9 (1) | <0.0001**** | 1.0 (1) | n.s. | 12.2 (1) | <0.0001**** |
| Capture time (baseline immobility subtracted) | 23.3 (1) | <0.0001**** | 0.9 (1) | n.s. | 16.3 (1) | <0.001*** |
| Time to 1st approach | 2.0 (1) | n.s. | 1.3 (1) | n.s. | 6.8 (1) | 0.009** |
| Time to 1st approach (baseline immobility subtracted) | 1.6 (1) | n.s. | 1.2 (1) | n.s. | 8.5 (1) | 0.004** |
| approach frequency (approaches/second) | 20.1 (1) | <0.0001**** | 4.0 (1) | 0.047* | 3.7 (1) | 0.053 |
| p (contact \| approach) | 15.7 (1) | <0.0001**** | 0.8 (1) | n.s. | 1.0 (1) | n.s. |
| p (capture \| contact) | 11.6 (1) | 0.0006*** | 2.7 (1) | n.s. | 6.7 (1) | 0.01 ** |
| Mean contact duration | 2.9 (1) | 0.09 | 4.2 (1) | 0.04* | 2.0 (1) | n.s. |
| Total contact duration | 16.2 (1) | <0.0001**** | 13.2 (1) | 0.0003*** | 22.0 (1) | <0.0001**** |
| Baseline immobility | 10.5 (1) | 0.0001**** | 0.5 (1) | n.s. | 1.2 (1) | n.s. |
| Arrests during approach | 0.26 (1) | n.s. | 0.002 (1) | n.s. | 1.4 (1) | n.s. |
| Baseline distance tr. | 25.4 (1) | <0.0001**** | 0.89 (1) | n.s. | 18.0 (1) | <0.0001**** |
| Approach distance tr. | 18.1 (1) | <0.0001**** | 2.4 (1) | n.s. | 3.9 (1) | 0.048* |
| Baseline speed | 3.7 (1) | 0.053 | 0.06 (1) | n.s. | 1.1 (1) | n.s. |
| Approach speed | 14.1 (1) | 0.0002*** | 0.2 (1) | n.s. | 1.1 (1) | n.s. |
| Abs. prey azimuth | 5.9 (1) | 0.01* | 8.6 (1) | 0.003** | 4.0 (1) | 0.044* |

### PSD-95 KO mice exhibit deficits in diverse epochs of hunting but partially improve with monocular vision

The hunting behaviour of mice is comprised of sequential epochs starting from initial prey detection, approaching and ultimately, contacting and capturing the prey using a combination of grasping and biting sequences^12–14^. Ipsilaterally projecting retinal ganglion cells (ipsi-RGCs) have previously been shown to signal prey-related visual cues^14^. Moreover, monocular vision is known to cause deficits in different phases of predatory behaviour such as prey detection, approach, contact and capture behaviours in adult WT mice^14^. Therefore, we set out to test the outcome of monocular vision in PSD-95 KO mice with respect to the approach, contact and capture epochs, in addition to capture time. Approach phases towards prey can be quantitatively defined by a combination of mouse speed, the mouse-cricket distance, the azimuth with respect to its prey and how these parameters change over time (*Figure 3A*).

**Figure 3.**
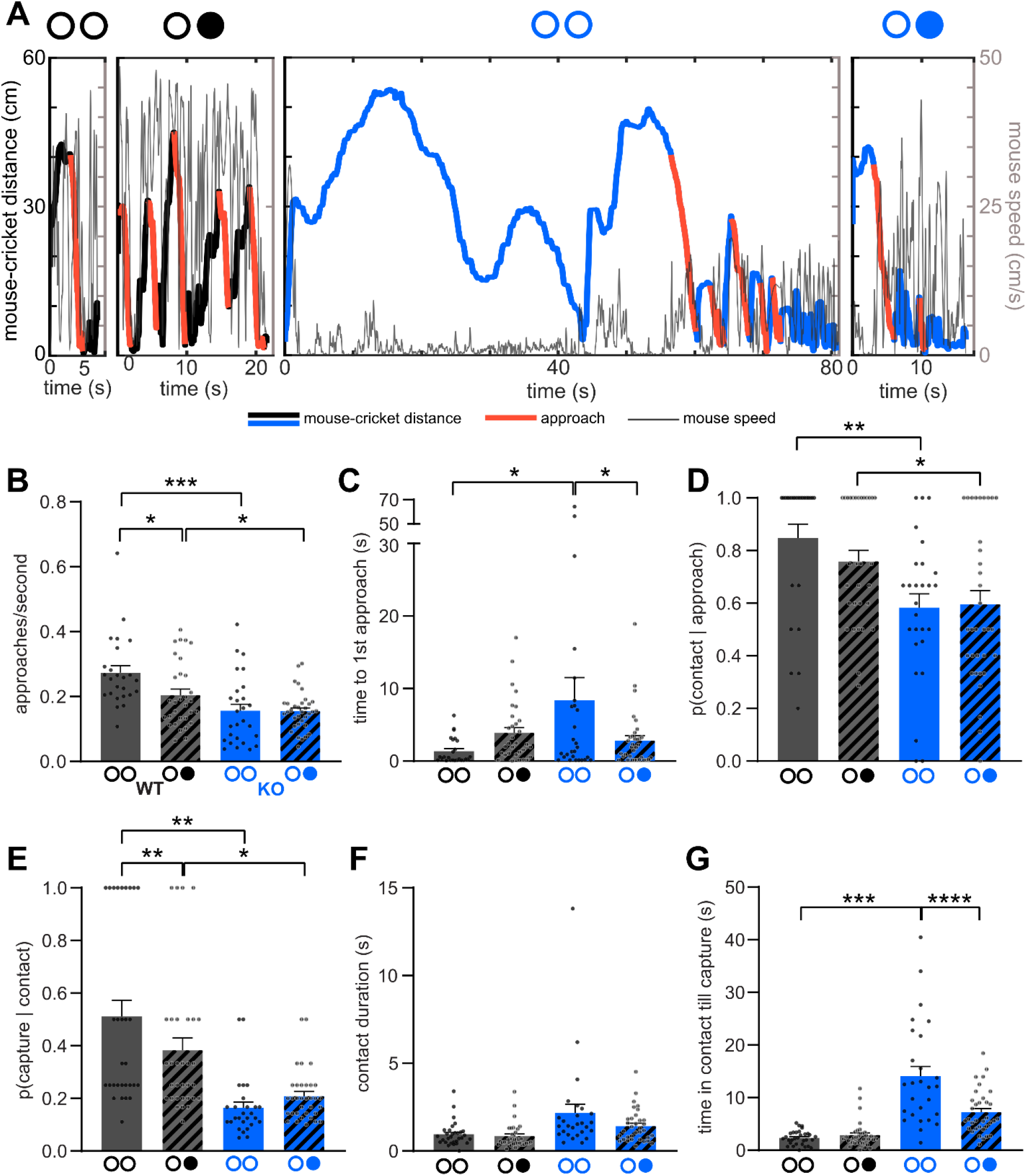
Binocular PSD-95 KO mice exhibit deficits in diverse epochs of prey capture behaviour but partially improve with monocular vision. (A) Example traces of mouse-cricket distance and speed over time until successful capture for an individual WT (black lines, left two panels) and KO (blue lines, right two panels) mouse with binocular and monocular vision, respectively. Approach epochs are labelled in red. (B) Rate of approaches per second towards prey. WT binocular: 0.27±0.02, n=25; WT monocular: 0.2±0.02, n=31; KO binocular: 0.16±0.02, n=27; KO monocular: 0.15±0.01, n=34. (C) Latency to initiate the first approach sequence towards prey. WT binocular: 1.38±0.35 s, n=25; WT monocular: 3.89±0.75 s, n=31; KO binocular: 8.37±3.13 s, n=27; KO monocular: 2.84±0.65 s, n=34. (D) Probability of a contact given a successful approach. WT binocular: 0.85±0.05, n=25; WT monocular: 0.76±0.04, n=31; KO binocular: 0.58±0.05, n=27; KO monocular: 0.6±0.05, n=34. (E) Probability of capturing prey given a successful contact. WT binocular: 0.51±0.06, n=30; WT monocular: 0.38±0.05, n=32; KO binocular: 0.17±0.02, n=27; KO monocular: 0.21±0.02, n=34. (F) Duration of each contact with prey. WT binocular: 1±0.12 s, n=31; WT monocular: 0.85±0.12 s, n=32; KO binocular: 2.17±0.5 s, n=27; KO monocular: 1.42±0.16 s, n=34. (G) Total duration of contact with prey. WT binocular: 2.35±0.24 s, n=31; WT monocular: 2.88±0.45 s, n=32; KO binocular: 14.09±1.87 s, n=27; KO monocular: 7.24±0.72 s, n=34. Bars represent mean±SEM. Each data point corresponds to a trial. Data points in E and F correspond to the mean value in a single trial. Bars of WT and KO mice are indicated by black and blue, respectively. ○○, binocular ○●, monocular. Linear mixed effects model with estimated marginal means for multiple comparisons (see Table 1 and S1). *p<0.05, **p<0.01, ***p<0.001 and ****p<0.0001.

WT mice generally approached prey more frequently compared to KO mice with both binocular and monocular vision (*Figure 3B*). The frequency of approaches decreased for WT mice with monocular vision, suggesting poorer prey recognition under monocular conditions. In contrast, in KO mice, both binocular and monocular performance displayed similar rates of approaches, which were reduced compared to WT mice. These results suggest that prey recognition was impaired in KO mice. With monocular vision, the visual field is reduced and resulted in a reduction of approaches in the WT mice. In contrast, in PSD-95 KO mice, closing an eye did not reduce approach frequency, indicating that despite the reduced visual field under monocular vision, something improved during monocular vision in the KO mice, compensating for the reduced visual field.

With binocular vision, KO mice were ∼6x slower compared to WT mice in detecting and initiating the first approach towards the cricket (*Figure 3C*), further supporting the impaired prey recognition under binocular vision in PSD-95 KO mice. Again, with monocular vision, KO mice, unlike WTs, were ∼3x faster to detect and initiate approaches towards the prey than with binocular vision. Together, these results indicate that binocular vision is severely compromised in PSD-95 KO mice. In contrast, monocular vision minimally affects predation and can even improve it compared to binocular vision in KO mice.

In addition, PSD-95 KO mice, with binocular vision, exhibited a lower probability than WT mice to successfully contact the crickets (mouse-cricket distance < 3 cm) once an approach sequence was initiated (*Figure 3D*) and a lower capture probability given a successful contact (*Figure 3E*). Both contact and capture probabilities did not change with monocular vision, unlike in WT mice where there was a significant reduction in their capture probability, but not in contact probability. Since PSD-95 KO mice did not improve with monocular vision, the reduction in PSD-95 KO mice suggests that these features were not solely dependent on binocular vision and thus additional prey capture impairments of PSD-95 KO mice may exist.

One characteristic feature of prey capture efficiency is how much time predators investigate or tend to spend in contact with their prey until successful capture. Compared to mice experienced in hunting, both juvenile and monocularly enucleated adult mice spend a much longer time in proximate contact with prey^14,18^. Although both genotypes, using both binocular and monocular vision, spent similar amounts of time in each individual contact with the crickets (*Figure 3F*), the total time spent in proximate contact with crickets in each trial with intact binocular vision was on an average ∼7x longer in KO compared to WT mice (*Figure 3G*). Notably, using monocular vision, KO mice exhibited an almost twofold reduction in the total proximate contact duration with prey until capture, i.e., strongly improved capture efficiency. Monocular vision did not exert any significant effect on the total prey contact duration for WT mice, contrary to what was previously reported in the literature^14^, most likely due to methodological differences in the nature of visual deprivation (eye closure vs. enucleation).

Thus, in comparison to WT mice, KO mice with binocular vision, were found to exhibit deficits in almost all (5/6) prey pursuit related parameters; with monocular vision, PSD-95 KO mice improved in two of these parameters, whereas WT mice deteriorated in two others (*Figure 3B-G*). Altogether, our analyses of approach, contact and capture epochs suggest that PSD-95 KO mice have profound deficits in hunting behaviour and that monocular vision improves hunting outcomes in the same KO animals. Our data also recapitulate the prevalence of deficits in the hunting behaviour of WT mice using monocular vision.

### Motor deficits of PSD-95 KO mice were partly due to binocular vision impairments

Since PSD-95 function might also be involved in maintaining optimal motor coordination^42^ and is associated with neurodevelopmental disorders, such as autism-spectrum disorders (ASD) or schizophrenia^43–45^, we set out to test whether the observed deficits of PSD-95 KO mice in hunting behaviour can be attributed to differences in the modulation of locomotion and/or their immobility levels when exposed to crickets. To establish whether the reduced hunting capabilities were the result of general motor deficits, we quantified travelled distance, speed and immobility/arrest-like states of the experimental animals during both non-approach (baseline) and approach phases. With binocular – but not monocular – vision, KO mice spent 4x more time immobile (mouse speed < 0.05 cm/s during non-approach phases) at baseline, compared to WT mice (*Figure 4A*). On the other hand, no differences were observed in baseline immobility between binocular and monocular vision for both genotypes. The percentage of arrest-like states during approach (2cm/s < mouse speed > 0.5cm/s) was neither influenced by genotype nor by visual condition (*Figure 4B*). To validate whether the overall duration of immobile phases impacted our quantified data, we subtracted the time spent immobile from the measured capture and detection times (*Figure S2*) and found that the observed differences remained. This suggests that the monocular improvements in KO mice are likely not influenced by their diminished immobility.

**Figure 4.**
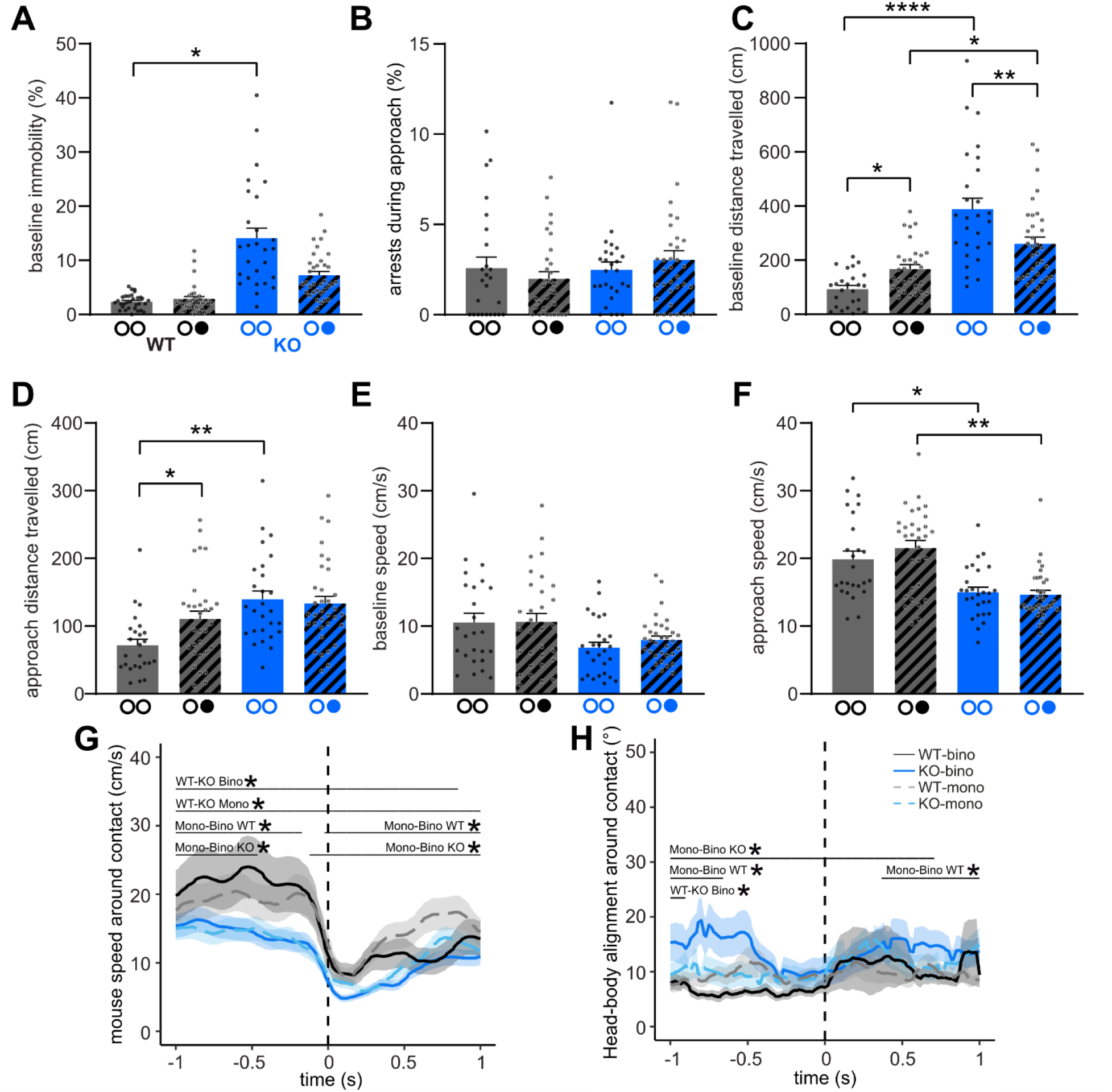
PSD-95 KO mice exhibit inefficient appetitive locomotion but improve with monocular vision. (A) Percentage of time mice stay immobile at baseline (non-approach phases). WT binocular: 2.35±0.24%, n=31; WT monocular: 2.88±0.46%, n=32; KO binocular: 14.09±1.87%, n=27; KO monocular: 7.24±0.72%, n=34. (B) Percentage of time mice are in an arrest-like state during an active approach towards prey. WT binocular: 2.58±0.61%, n=25; WT monocular: 1.99±0.38%, n=31; KO binocular: 2.48±0.44%, n=27; KO monocular: 3.04±0.5%, n=34. (C) Distance travelled at baseline when mice are not actively approaching prey. WT binocular: 92.96±12.99 cm, n=25; WT monocular: 167.3±16.19 cm, n=31; KO binocular: 388.5±40.25 cm, n=27; KO monocular: 260.6±24.8 cm, n=34. (D) Distance travelled during approach epochs towards prey. WT binocular: 71.74±8.92 cm, n=25; WT monocular: 110.5±11.68 cm, n=31; KO binocular: 139.4±12.39 cm, n=27; KO monocular: 133.2±10.68 cm, n=34. (E) Speed of mice at baseline (non-approach phases). WT binocular: 10.54±1.36 cm/s, n=25; WT monocular: 10.65±1.24 cm/s, n=31; KO binocular: 6.81±0.8 cm/s, n=27; KO monocular: 7.97±0.59 cm/s, n=34. (F) Speed of mice during the active approach phases. WT binocular: 19.86±1.22 cm/s, n=25; WT monocular: 21.55±1.1 cm/s, n=31; KO binocular: 15.01±0.73 cm/s, n=27; KO monocular: 14.64±0.63 cm/s, n=34. (G) Speed of mice around ±1 s of contact with prey. Lines and shaded areas indicate mean±SEM. Dashed line indicates the time of a contact. (H) Head-to-body alignment of mice around ±1 s of contact with prey, with body and head completely aligned at an angle of 0. Lines and shaded areas indicate mean±SEM. Dashed line indicates the time of a contact. Bars indicate mean±SEM. Each data point in A and B corresponds to % of immobility in each trial. Data points in E and F correspond to the median mouse speed in a trial. ○○, binocular ○●, monocular. Linear mixed effects model with estimated marginal means for multiple comparisons (see Table S1). *p<0.05, ***p<0.001.

The degree of appetitive locomotion during hunting^46^ and how it changes in different sensory conditions can be assessed by quantifying how much distance mice covered during baseline (non-approach) phases (*Figure 4C*) and active prey approach phases (*Figure 4D*); reduced appetitive locomotion will lead to significantly more distance traversed during baseline vs. approach phases, possibly due to impaired prey recognition. Indeed, in WT mice, the traversed distances did not differ between trial-matched baseline and approach phases with binocular vision (baseline: 92.96±12.99 cm, approach: 71.74±8.92 cm; Wilcoxon’s test, *p*=0.15). However, with monocular vision, the traversed distances were significantly higher during baseline vs. approach phases (baseline: 167.3±16.19 cm, approach: 110.5±11.68 cm; *p*<0.0001), thus indicating reduced relative appetitive locomotion in WT mice with monocular vision. In contrast, KO mice exhibited significantly reduced relative appetitive locomotion with both binocular (baseline: 388.5±40.25 cm, approach: 139.4±12.39 cm; *p*< 0.0001) and monocular vision (baseline: 260.6±24.8 cm, approach: 133.2±10.68 cm; *p*<0.0001). The distance traversed at baseline (*Figure 4C*) by KO mice when not in active pursuit was ∼4x higher compared to WT mice with binocular vision and more than 2x higher with monocular vision. Markedly, WT mice traversed more than 2x the distance during baseline using monocular compared to binocular vision, whereas KO mice did the opposite: they covered ∼33% less distance with monocular vs. binocular vision. During approaches towards prey with binocular vision, KO mice also covered ∼2x the distance compared to WT mice (*Figure 4D*). However, monocular vision did not affect the distance KO mice covered during approaches towards prey whereas WT mice travelled significantly higher distances. Overall, our data indicate that in PSD-95 KO mice, contrary to WT mice, monocular vision facilitates a relative improvement in appetitive locomotion, despite a general deficit in locomotion.

While both WT and KO mice maintained similar average speeds when not actively approaching crickets (*Figure 4E*), KO mice were generally slower compared to their WT counterparts during approach phases (*Figure 4F*), both binocularly (WT: 19.86±1.22 cm/s, KO: 15.01±0.73 cm/s) and monocularly (WT: 21.55±1.1 cm/s, KO: 14.64±0.63 cm/s). This was complemented with a generally slower mouse speed right before individual proximal contacts with prey in the KOs (*Figure 4G;* significant WT-KO Bino difference from −1 to 0.9 s and WT-KO Mono difference from −1 to 1 s), thereby underscoring a genotype difference in speed modulation before contacting prey. To investigate orienting movements, associated with head movements, we quantified the alignment between the mouse’s head and body^15^. This showed a difference between binocular and monocular head-body alignment around contact (*Figure 4H*), for KO mice there is an increased head-to-body angle with binocular vision from the 1 s leading up to contact until 0.75 s after contact, whereas for WT mice there is a decreased head-to-body angle with binocular vision from −1 to - 0.65 s and 0.4 to 1 s. Mouse speed and head-to-body alignment in the 2.5 s leading up to contact have been depicted in *(Figure S4)*.

Taken together, our results indicate that the observed and quantified deficits in predatory behaviour of PSD-95 KO compared to WT mice are not due to higher levels of immobility in the KO mice. The fact that with monocular vision, KO mice travelled shorter distances than with binocular vision when not actively pursuing prey, despite lower speeds during prey approaches under both visual conditions, points towards improved efficiency of KO mice in regulating appetitive locomotion with monocular vision.

### PSD-95 KO mice exhibit binocular visual field bias with monocular vision

During active prey pursuit, adult WT mice are known to dynamically orient their head position (prey azimuth) to keep their prey within the binocular visual field (∼40°), a characteristic termed the “binocular visual field bias”^13,14,18^. We wanted to know whether the deficits in predation displayed by PSD-95 KO mice resulted from a lack of binocular visual field bias. WT mice exhibited a strong binocular visual field bias during approach phases (with binocular vision) before contact with prey (*Figure 5*), as has been previously reported^13,14^. Notably, KO mice, irrespective of monocular or binocular vision, exhibited a WT-like progressive decrease in prey azimuth as a function of the mouse-cricket distance (*Figure 5A*). Moreover, at the end of active approaches towards the crickets, both binocular WT and KO mice maintain prey azimuth within the binocular zone. In turn, the same WT animals, with monocular vision, have a ∼3-fold decrease in the probability of the azimuth being in the binocular zone (*Figure 5B*). In contrast, KOs, with both binocular and monocular vision, maintained prey azimuth within the binocular visual field. Thus, despite the lateral location of the eye at the side, they maintained the prey in the binocular visual field. Immediately following contact, the azimuth increases for all mice (*Figure 5C*). Although the azimuth of KO mice with binocular vision does not increase as starkly as WT (not significant), it is substantially lower compared to monocular vision (significantly different between −0.15 to 1s). Additionally, there was a significant effect of monocular vision for both WT (significant between −0.9 to 1s) and KO mice (−1 to −0.6s). Finally, at the start of the approach epoch, KO mice with monocular vision had a significantly higher azimuth than WT mice with monocular vision (−1 to −0.75s). At the end of the approach epochs, the mean azimuth was similar between WT and KO mice with binocular vision but with monocular vision, the azimuth was ∼33% higher in WT mice, whereas it remained similar in KO mice (*Figure 5D*). Mouse azimuth in the 2.5 s leading up to contact has been depicted in *(Figure S4)*. Reiterating the result above, whereas the WT azimuth on average was ∼2-fold higher with monocular vision than with binocular vision, no significant change was observed in the KO mice. Thus, the absence of PSD-95 did not disrupt the binocular field bias for either visual condition, suggesting that monocular visual cues are sufficient to preserve this bias for the KO mice.

**Figure 5.**
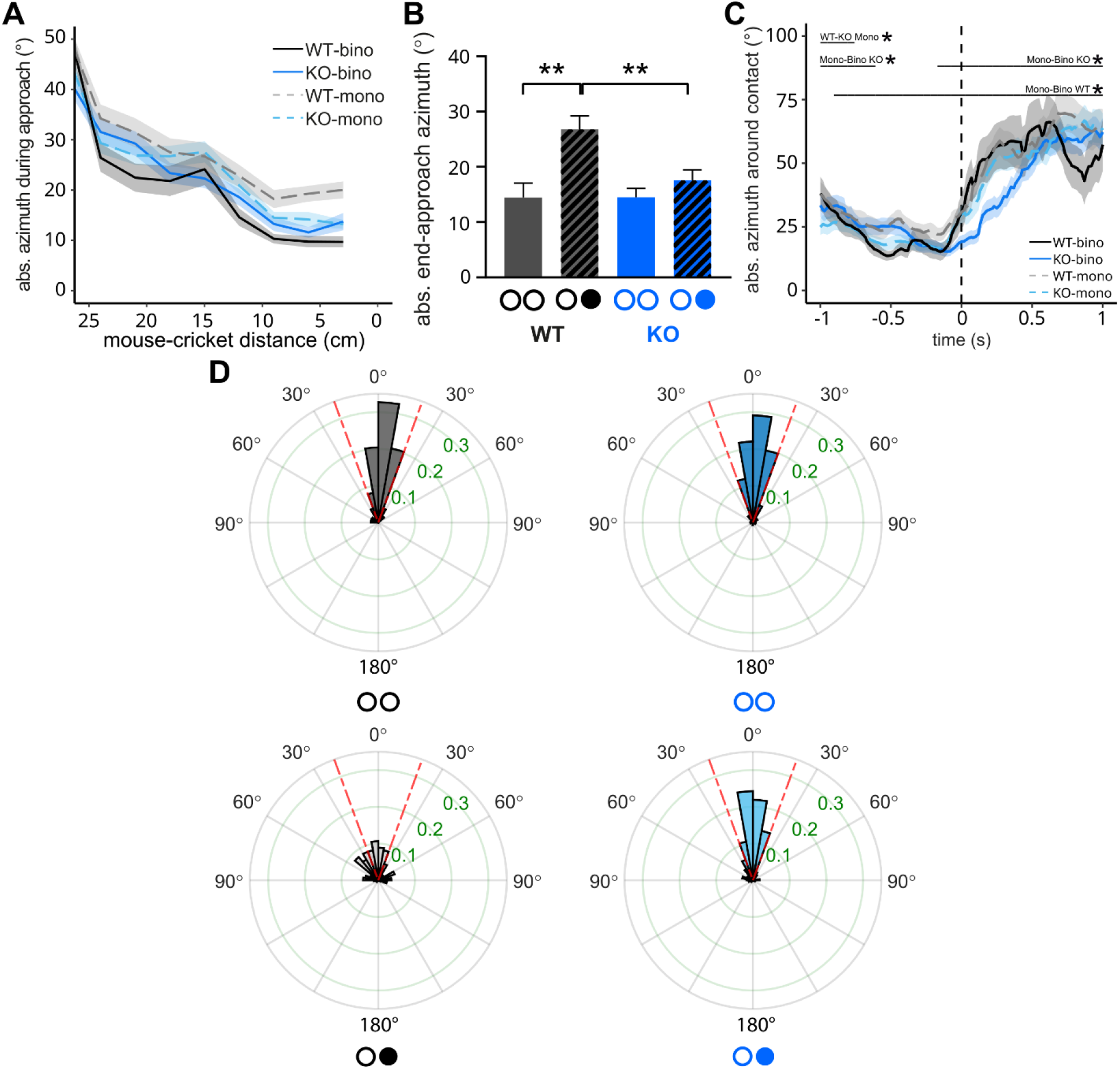
Binocular visual field bias is conserved in KO mice irrespective of visual condition (binocular/monocular), unlike in WT mice. (A) Mean absolute azimuth as a function of mouse-cricket distance until contact during approach phases. (B) Polar probability histograms (10° bins) of azimuth at the end of approach towards prey. Dashed red line indicates the binocular visual field (±20°). (C) Mean absolute azimuth ±1 s of contact with prey. Lines and shaded areas indicate mean±SEM. Dashed line indicates the time of a contact. (D) Absolute azimuth at the end of approach towards prey. WT binocular: 14.4±2.62°, n=49; WT monocular: 26.8±2.42°, n=97; KO binocular: 14.5±1.61°, n=140; KO monocular: 17.6±1.9°, n=148. Bars indicate mean±SEM. ○○, binocular ○●, monocular. Linear mixed effects model with estimated marginal means for multiple comparisons (see Table 1 and S1). *p<0.05, **p<0.01.

### WT and KO mice display similar disruptions of predation under darkness

The critical role of vision for the animals’ predatory behaviour was demonstrated by comparing performance between light and dark trials^13^. Under darkness, predatory behaviour of adult WT mice was previously shown to be severely compromised: without perceptible visual cues, WT mice have deficits in prey detection, recognition, approach and subsequent capture, and fail to maintain prey within the binocular visual field^13^. Furthermore, deficits in predation observed with monocular vision are absent under darkness^16^, confirming the importance of visual processing for successful predation. To assess whether the genotype differences we observed under binocular/monocular conditions were indeed dependent on vision, additional prey capture trials were conducted under complete darkness. No significant differences were observed between genotypes or eye conditions during darkness (*see Table S2*) for capture times (*Figure S3A*), approach frequency (*Figure S3B*), prey detection times (*Figure S3C*), contact and capture probabilities (*Figure S3D-E*) and total prey contact duration (*Figure S3F*). WT mice were slightly more immobile during baseline with monocular vs. binocular vision while KO mice exhibited similar levels of immobility between eye conditions (*Figure S3G*). KO mice, contrary to their WT counterparts, traversed greater distances during both baseline (*Figure S3H*) and approach phases with both bino- and monocular vision in the dark (*Figure S3I*). Moreover, contrary to observations under normal illumination conditions, there was no perceptible difference between bino- and monocular vision in terms of locomotor behaviour during baseline and approach epochs in KO mice under darkness. This indicates that the monocular improvement in relative appetitive locomotion in KO mice is also vision dependent. WT and KO mice also maintained similar speeds at baseline (*Figure S3J*) and when actively approaching crickets (*Figure S3K*). Furthermore, both genotypes with binocular and monocular vision, failed to maintain the prey azimuth within their binocular visual field under darkness (*Figure S3L-N*). In summary, the lack of any perceptible differences between binocular and monocular vision within and between genotypes in darkness, strongly indicate that the observed deficits and the improvements in predation observed with monocular vision in PSD-95 WT and KO mice, respectively, are primarily vision dependent, and thus the result of experience-dependent changes in the visual processing machinery.

### KO mice are better at orientation discrimination with monocular vision

PSD-95 KO mice are poorer at orientation discrimination using binocular vision compared to WT mice despite WT-like visual acuity^34^ in the VWT^33^ (*Figure 6A*). Since CP for OD-plasticity does not close in PSD-95 KO mice^32^ and binocular matching occurs during CP^25^, we reasoned that the observed deficit in orientation discrimination of PSD-95 KOs might be due to an interfering, somehow non-matched input from the other eye, and that performance might improve under monocular vision, as shown for prey capture. We therefore assessed whether monocular vision would change performance in both WT and PSD-95 KO mice. PSD-95 KO mice took significantly longer than WT mice to learn the task (i.e., reach 70% accuracy at the lowest possible orientation difference) and enter the test phase (*Figure 6B;* χ²(1)=25.7, p<0.0001***). During the test phase, we determined the smallest orientation difference that could be discriminated with at least 70% accuracy (*Figure 6C*) for both genotypes and visual conditions. Using binocular vision, WT mice needed an orientation difference of at least ∼15° to discriminate sine wave gratings, which increased ∼2-fold using monocular vision. This finding underscores the importance of binocular visual processing for optimal orientation discrimination in mice which was previously described in humans^47,48^. In contrast, the performance of PSD-95 KO mice strongly improved using monocular vision: While KO mice displayed a pronounced orientation discrimination deficit with binocular vision, and needed an orientation contrast of ∼52°, and thus ∼3 times higher than WT mice (∼17°, *p*<0.0001), their monocular performance (∼32°) was indistinguishable from monocular WT mice (∼34°, *p*=0.79). The monocular improvement in orientation discrimination in PSD-95 KO mice thus corroborates with the improvements observed in the prey capture task.

**Figure 6.**
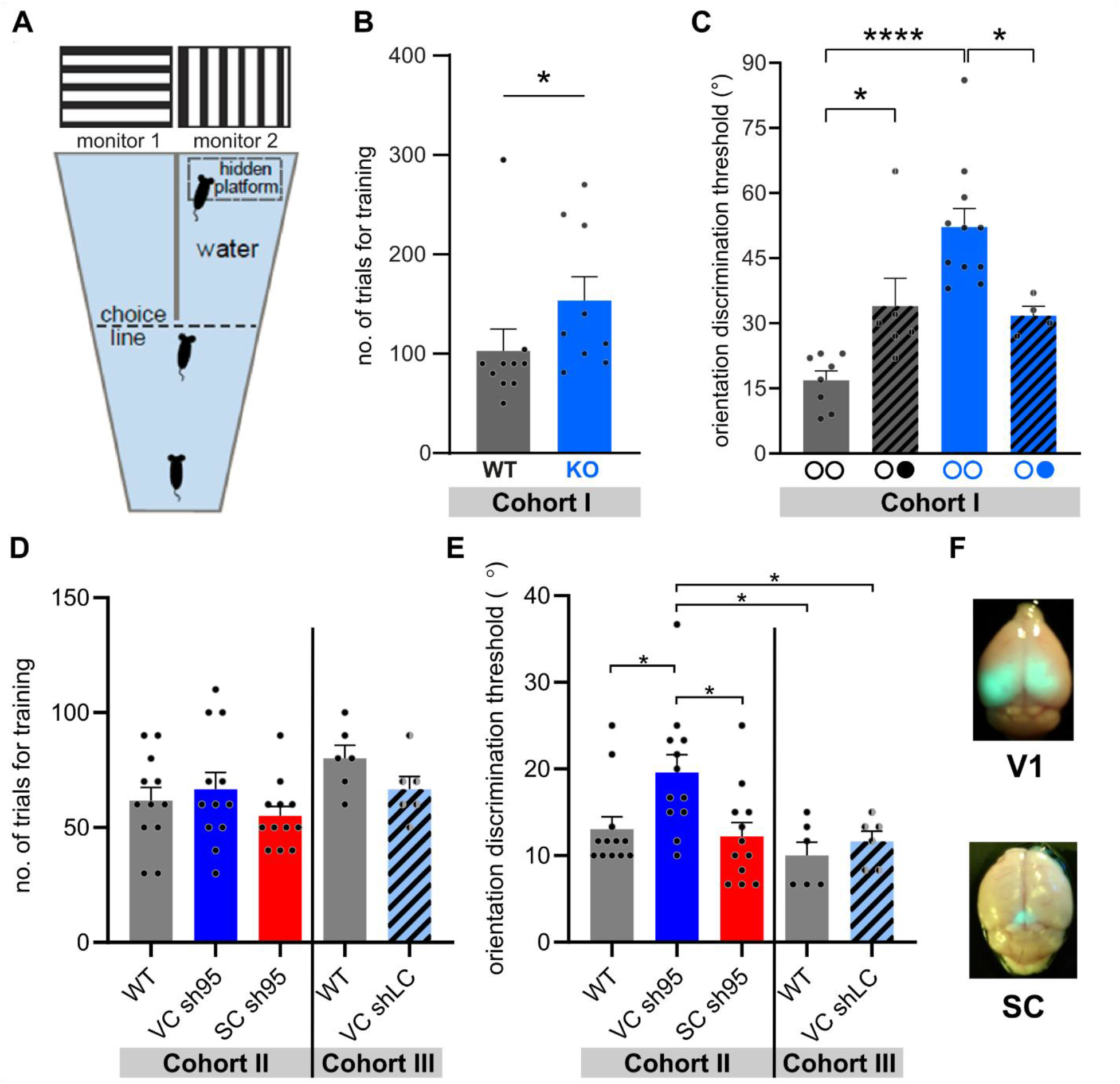
PSD-95 KO mice exhibit poorer orientation discrimination compared to WT mice but improve monocularly. (A) Schematic of the visual water task adapted with permission for testing orientation discrimination threshold (Favaro et al., 2018). (B) Total number of trials required for training WT and KO mice before beginning the test phase. WT: 103±24, n=10; KO: 153±24, n=9; Mann-Whitney, p=0.016. (C) Orientation discrimination thresholds of WTs and KOs with binocular and monocular vision during the test phase. WT binocular: 17±2°, n=8; WT monocular: 34±6°, n=6; KO binocular: 52±4°, n=11; KO monocular: 32±2°, n=4. Multiple linear regression with estimated marginal means for multiple comparisons. F_3,25_=14.17, *p*<0.0001; β_g_= −35.45, *p*<0.0001; β_v_=-20.52, *p*=0.0065; β_g*v_=37.53, *p*=0.0005. WT_b_ - WT_m_: t(25)=2.66, *p*=0.018; KO_b_ - KO_m_: t(25)=2.97, *p*=0.013; WT_b_ – KO_b_: t(25)=6.45, *p*<0.0001; WT_m_ - KO_m_: t(25)=0.273, p=0.787. (D) Total number of trials required for training WT cohort 2 and 3, VC sh95KD, SC sh95KD, and VC shLC mice before beginning the test phase. WT cohort 2 (left): 62±20, n=12; VC sh95 KD 67±25, n=12; SC sh95 KD 55±15, n=12; WT cohort 3 (right): 80±14, n=6; VC shLC 67±14, n=6. Multiple linear regression with estimated marginal means for multiple comparisons. F_4_=1.87, p=0.13; β_g_=61.67, p<0.0001. (E) Orientation discrimination thresholds of WT cohort 2 and WT cohort 3, VC sh95KD, SC sh95KD, and VC shLC during the test phase. WT cohort 2 (left): 13±5°, n=12; VC sh95 KD 20±7°, n=12; SC sh95 KD 12±6°, n=12; WT cohort 3 (right): 10±4°, n=6; VC shLC 12±3°, n=6. Multiple linear regression with estimated marginal means for multiple comparisons. F_4_=4.61, p=0.003; β_g_=13.06, p<0.0001. WT1 – VC sh95: t(43)=-2.93, p=0.01; WT2 – VC sh95: t(43)=-3.50, p=0.01; VC shLC – VC sh95: t(43)=-2.90, p=0.01; VC sh95 – SC sh95: t(43)=3.30, p=0.01. (F) Representative examples of viral expression in V1 or SC, visualised with a blue light torch and an orange filter. Bars indicate mean±SEM. ○○, binocular ○●, monocular. *p<0.05, ****p<0.0001.

We next investigated the differential contribution of PSD-95 expression in V1 and SC for orientation discrimination. Whereas V1 is, among other processes, involved in encoding binocular information, SC is implicated in numerous predation-relevant visual processing^18,24,49–51^. Moreover, PSD-95 has been shown to mediate SC synaptic maturation^52^. In cohort II, mice either received a knock-down of PSD-95 in V1 (sh95 V1) or SC (sh95 SC) or no intervention at all (WT) and underwent the same orientation discrimination procedure as before. In cohort III, a second WT group was compared with mice who received a sham virus injection in V1 (shLC). No significant difference between the two WT groups were found. Moreover, in contrast to the PSD-95 KO, neither the V1 nor SC PSD-95 KD mice took significantly longer to learn the task (i.e., ≥90% accuracy on two consecutive sessions while completing ≥3 sessions). On the other hand, there was a significantly impaired orientation discrimination for mice with a V1-specific PSD-95 KD (sh95 V1; *Figure 6D*) compared to all other groups, including WT mice and animals with control virus injection (shLC V1). Notably, mice with a PSD-95 KD in SC (sh95 SC) exhibited normal orientation discrimination, supporting a role for PSD-95 expression in V1 in binocular integration.

Taken together, our results from both the behaviourally relevant prey capture task and from testing orientation discrimination using the VWT indicate that PSD-95 KO mice suffer from compromised binocular integration of visual inputs. Thus, in contrast to WT mice that always get worse using monocular vision, KO animals can even display improved visual performance using monocular vision.

## DISCUSSION

This study explored whether loss of PSD-95, associated with lifelong CP, influences binocular visual behaviour. To this end, we performed both monocular and binocular tests in the visual water task for orientation discrimination and in the ethologically relevant prey capture task. Notably, PSD-95 KO mice exhibited impaired binocular visual behaviour in both predation (*Figure 1-5*), and orientation discrimination (*Figure 6*) and strongly improved with monocular vision to performance comparable to WT mice in both tests. Importantly, impaired orientation discrimination was solely observed in mice with V1-specific knock-down of PSD-95 and not after knock-down in the superior colliculus (*Figure 6*).

The full range of functions served by binocular vision in mice and how it develops during CP remains elusive, partly because critical features of foveated species such as binocular summation are incompletely characterised and features of binocular integration were not pinned to critical period development. Moreover, the development of binocularity is mediated by experience-dependent maturation of excitatory cortical circuits^22,26,53^ and facilitated by the heightened plasticity during visual CP in V1^54,55^. In WT mice, binocular orientation discrimination was ∼2x better than the monocular performance and may therefore be similar to what is typically observed in humans for binocular summation^56^, the perceptual improvement when using binocular instead of monocular vision in tasks not requiring depth perception. The increase of binocular vision compared to monocular vision in mice depicts similarities in visual features with foveated species despite anatomical differences.

Strikingly, in PSD-95 KO mice we did not observe binocular improvement, but rather the opposite: both predation (*Figure 1-5*) and orientation discrimination (*Figure 6*) improved with *monocular* vision, and monocular performance in KO mice was similar to monocularly tested WT mice. This binocular deficit was apparent from training onset, as no PSD-95 KO mice caught a cricket on the first training day. Additionally, PSD-95 KO mice took longer to reliably catch crickets and required more trials for training on the VWT. A decrease or absence of PSD-95 has been linked to learning deficits, e.g. impaired memory retention^57,58^, impaired recognition memory and learning the T-maze^59^, as well as impaired conditioned place preference^60,61^. This impaired learning was specific to the PSD-95 KO mice, with neither a local knock-down in the SC nor V1 resulting in similar learning impairments. The PSD-95 mediated learning effect may therefore be attributed to regions typically linked to learning of sensory information. For instance, local PSD-95 knock-down in dorsal CA1 of the hippocampus impaired training-induced accuracy of spatial choice, quantified by the number of visits to a rewarded location in young adult mice^62^. Similarly, KD of PSD-95 in the rat gustatory cortex impaired learning of novel tastes^63^. In contrast to learning deficits, which may contribute to the observed slower learning rate, disrupted binocular integration is likely to result in the observed binocular and not monocular predation impairment, supported by similar learning trajectories in the two visual conditions. Improved visual perception with monocular vs. binocular vision under neurotypical conditions is rarely observed in humans^56,64–67^. It can typically be observed with amblyopia, where the visual acuity between both eyes differ^68,69^. However, a selective amblyopia with PSD-95 KO is highly unlikely, since visual acuity of both eyes of PSD-95 KO mice did not differ^32^, suggesting alternative causes for the binocular impairment.

Mice are known to utilise monocular cues in nature: for instance, using the ethologically relevant visual gap crossing task, it was reported that mice can estimate distance cues using motion parallax even with monocular vision^70^. The improvements observed with monocular vision in the KO mice might be a consequence of improved visual capabilities due to the elimination of the binocular interference caused by the non-matched inputs from either eye. This supports the idea that the KO mice, with monocular vision, may more efficiently utilise monocular cues such as motion parallax, relative object size and height, instead of using binocular visual information like depth and disparity cues, compared to WT mice. Strikingly, visual acuity (measured with VWT), which is physiologically also dependent on CP and visual-experience^71^, is unaffected in PSD-95 KO mice^34^. This is likely explained by our previous observation that PSD-95 KO does not alter the maturation of local inhibitory networks in V1^32,72^, which are required for visual acuity development^73^. Notably, driving the maturation of inhibitory networks with BDNF, even in the absence of experience, can rescue visual features including visual acuity^74,75^. Thus, the phenotype of PSD-95 KO mice is primarily driven by impaired excitatory network refinement.

Consistent with the absence of binocular improvement observed in predation, PSD-95 KO mice exhibited a comparable modulation of orientation discrimination in the VWT (*Figure 6*): PSD-95 KO animals could distinguish smaller orientation differences, i.e., improved orientation discrimination performance with monocular vision. These data are consistent with a mismatch in binocular integration that may lead to binocular interference. Notably, an orientation discrimination deficit was solely observed following PSD-95 knock-down in V1 but not in SC. The monocular improvement in predation of PSD-95 KO mice indicates the potential role of binocular integration, and consequently depth perception^76^ in prey capture behaviour, suggesting a role of PSD-95 in binocular integration and depth perception. However, orientation discrimination measured with VWT relies on visual perception and binocular integration, not depth perception^47,48^, thus, supporting a pivotal role of PSD-95 in binocular integration. This finding is supported by clinical studies which report enhanced monocular performance of strabismic patients in orientation discrimination^77^ and improvements in visual perception of adult amblyopes after monocular perceptual learning^78,79^. Individuals with ASD exhibit impaired binocular integration or increased binocular interference^80–82^, and strikingly, the Williams syndrome, an ASD-related syndrome associated with PSD-95^42,83^, exhibited impaired orientation discrimination^84^, echoing our findings. In contrast, our WT mice exhibited binocular improvements reminiscent of binocular summation. Moreover, binocular summation is known for a starker impact at low stimulus contrasts and short durations^85,86^. Whether binocular summation in mice follows similar patterns as observed in humans should be tested in future studies. Here, we did not observe binocular, but monocular improvements for PSD-95 mice. We suspect that PSD-95 KO mice may not lack binocular summation, but instead suffer from binocular interference, as binocular summation has been observed clinically in both healthy individuals as well as patients with eye diseases^87^. Therefore, while a low stimulus contrast or signal duration may affect binocular summation observed in WT mice, the impact of low stimulus contrast or signal duration in PSD-95 KO mice remains unpredictable. Nonetheless, we predict that the binocular interference of PSD-95 KO mice persists and performance with monocular vision to remain superior to that with binocular vision even under these conditions.

In contrast to other predation parameters, we found that adult KO mice, with both binocular and monocular vision, maintain a strong binocular visual field bias, whereas WT mice fail to maintain this bias with monocular vision (*Figure 5*). Mice utilise their large binocular visual field during active prey pursuit^13,18^ by maintaining the prey image in the upper-temporal retinal field, an area that corresponds to the lower-central visual field^88^. Not keeping the prey within the binocular field impairs prey pursuit, as demonstrated with juvenile and monocularly deprived adult WT mice, who failed to keep prey efficiently within the binocular visual field^13,14,18^. The accurate orienting for sustained approaches towards prey – signified by the prey azimuth being within the binocular visual field and functionally attributed to narrow-field cells in the superior colliculus^89^ – was not affected in both binocular and monocular KO mice, contrary to observations in WT mice. While this may indicate PSD-95 independent compensatory mechanism(s), such as the optokinetic reflex, we hypothesise that a changed binocular field bias in WT mice with monocular vision results from mice optimising visual input. While humans close one eye to optimize visual input, mice are not known to close one eye to optimize vision as they are only known to squint during pain. Therefore, with monocular vision WT mice might adjust their visual field bias to optimize visual input, whereas PSD-95 KO mice would no longer experience binocular interference, thereby eliminating the need to optimize. Nonetheless, the mechanism through which PSD-95 KO but not WT mice with monocular vision retain appropriate field bias remain to be elucidated. Future studies could assess how receptive field tuning is affected in superior colliculus and bV1 neuronal populations following loss of PSD-95.

We propose that the images of the two eyes of the KO mice are not fused properly to generate a meaningful composite image^90^, resulting in binocular interference and visual impairments. Under this scenario, PSD-95-dependent maturation of AMPAR-silent synapses^32,38^ would be the crucial mechanism to refine excitatory networks for the proper fusion of images. In support, PSD-95 is required for the maturation of excitatory networks during CP^28,34,37,38,72^. Experience during CP is involved in the emergence of binocular vision-dependent processes, such as interocular matching of orientation preferences^26^, prey-capture learning^18,19^ and speed discrimination in V1 neurons^51^. This, in turn, likely interferes with the proper maturation and stabilisation of individual eye inputs onto binocular neurons, leading to non-matched binocular inputs onto single neurons^91^. How the matching of binocular inputs, such as orientation preference are affected in PSD-95 KO mice in comparison to e.g., proper maturation of inhibition will need further exploration. The brain regions in which PSD-95 asserts these visual effects have yet to be discovered. Nonetheless, V1, a central cortical hub for binocular integration across species^92–94^, is a region of interest due to its role in encoding orientation and direction tuning^20,21,95–99^ and phase disparity information^100–103^. Particularly in the context of predation, inactivation of V1 led to deficit in prey detection and capture times in adult mice^24^. Alongside V1, superior colliculus has been implicated in predation-relevant visual processing^18,24,49–51^. Recent studies have linked the emergence and refinement of predation during CP^18,19^, including in the context of ODP, with binocular neurons of SC^16^. More specifically, PSD-95 has been shown to mediate superior colliculus synaptic maturation by promoting juvenile superior colliculus LTP^52^. Both V1 inactivation^24^ and silencing superior colliculus wide-field neuron activity^89^ individually impair prey detection. Since prey detection is also impaired in PSD-95 KO mice with binocular vision, maturation of V1 and/or superior colliculus silent synapses might contribute to optimal prey detection with binocular vision. Our observation that orientation discrimination is impaired only after a V1-specific, but not superior colliculus-specific knock-down of PSD-95 strongly suggests a primary involvement of V1. Regardless, given the monocular improvement of prey detection in PSD-95 KO mice, it would be worthwhile to assess the role of PSD-95 for prey detection in binocular superior colliculus neurons.

To demonstrate the importance of vision in the PSD-95-dependent optimisation of hunting behaviour, we performed prey capture trials also in the dark, for both WT and KO mice. Our data reveal a lack of genotype and eye condition differences during darkness: both WT and KO mice exhibited a similar degree of impairment when having to hunt in darkness (*Figure S3*). This observation is in line with several previous studies; one reporting predation in adult WT mice to be compromised in darkness^13^, the second reporting compromised predation with monocular vision^14^, and the third exclusively reporting a predation deficit in monocular mice in the light, but not in the dark^16^. PSD-95 loss might also cause some sensorimotor impairments, as has been observed in predation of Shank3 KO mice, an ASD model^104,105^. Late-CP Shank3 KO mice exhibited heightened immobility both at baseline and approach towards prey, characterised by speed modulation deficits^19^. Reduced expression of PSD-95 has been implicated in hypersociability^106^, PFC-mediated learning and working memory deficits^59^, hippocampus-mediated spatial learning deficits^107^, and, potentially, in ASD and schizophrenia^44^. Indeed, we observed that even with sufficient training, PSD-95 KOs exhibited slightly elevated baseline immobility (*Figure 4*). On the other hand, during approach phases, WT and KO mice showed similar amounts of arrest-like states. Despite the slower speeds of KO compared to WT mice, their predation improvement with monocular vision is not likely caused by a differential modulation of speed: Speed modulation of KO mice was unchanged between binocular and monocular conditions under both light and dark conditions. KO mice travelled greater distances in comparison to WT mice, but with monocular vision, travelled shorter distances at baseline compared to WT. The improved efficiency in active prey pursuit and capture in KO mice might be a consequence of better prey detection and lower prey investigation times with monocular vision. Taken together, these observations allow us to conclude that the observed impairments/improvements in predation are primarily vision-dependent in both genotypes, and not due to differences in (appetitive) locomotion, nor caused by other sensory modalities.

The vision-dependent effects may also be affected by a difference in gaze or binocular coordination. However, binocular coordination is required for VWT, and VWT-derived visual acuity remained intact in PSD-95 KO mice ^34^. Gaze differences would be expected to also lower binocular visual acuity^108^, which was not observed in PSD-95 KO mice. Furthermore, a small gaze difference would prevent the summation effect of binocular vision and reduce orientation discrimination to the monocular threshold^109^. However, in PSD-95 KO mice, binocular orientation discrimination was worse than with monocular vision, consistent with our interpretation of an impaired binocular processing. Together with only subtle locomotion effects in the prey-capture task, impaired binocular integration resulting in binocular interference remains a likely explanation of the binocular and light-specific effects.

In summary, our results support an essential role of PSD-95-mediated maturation of AMPAR-silent synapses for optimal binocular visual processing, as shown for both the ethologically relevant prey capture task and for orientation discrimination in the VWT. Our results indicate compromised binocular integration of visual inputs in the visual circuity and that monocular vision can partly improve visually guided behaviour when binocularity is compromised due to the loss of PSD-95. We propose this to result from improper fusion of the individual images of the eyes, resulting in binocular interference and visual impairments. This underlines the importance of CP-mediated silent synapse maturation in optimising binocular vision-dependent ethologically relevant behaviours in mice, and furthermore, highlight similarities of binocular visual process between mice and humans.

### Limitations of the study

The main goal of this study was to assess the impact of PSD-95 mediated AMPAR-silent synapse maturation on naturalistic binocular visual behaviour in adult mice. Although, the behaviour as a whole was studied thoroughly, future studies should analyse to what extent eye movements might be affected due to PSD-95 loss. Furthermore, using region- and/or cell type-specific knock-down should reveal the role of PSD-95 for efficient predatory behaviour. Additionally, region-specific binocular integration in PSD-95 KD mice could be probed with electrophysiology or calcium imaging. To further support PSD-95 mediated CP closure impairing binocular predatory behaviour, future studies should compare prey-capture efficiency pre- and post CP closure. We also observed that female PSD-95 KO mice had massive problems to reliably hunt crickets: their prey capture success was either very low or highly inconsistent during the training phase, leading us to exclude 9/10 tested female KO mice from the current dataset. While investigating the underlying cause of this prominent sex difference is beyond the scope of the present study, it would be interesting to investigate a potential link between PSD-95 and sex differences in predation in the future.

## RESOURCE AVAILABILITY

### Lead contact

Further information and requests for resources should be directed to and will be fulfilled by the corresponding authors and the lead contact.

### Materials availability

No new materials or reagents have been generated in this study. However, further details about the behaviour arena can be obtained from the lead contact upon reasonable request.

### Data and code availability

Code and data generated and used in this study are available on a common public repository (https://doi.org/10.25625/O2OLFZ). Access to all original videos can be obtained from the corresponding authors/lead contact upon reasonable request.

## ACKNOWLEDGMENTS

Supported by DFG through the CRC 889 “Cellular Mechanisms of Sensory Processing” to SL (B5) and OMS (B3), and National Institutes of Health (MH131717 to OMS). We acknowledge support by the Open Access Publication Funds/transformative agreements of the Göttingen University. We thank Dr. Cris Niell and Dr. Jennifer Hoy for sharing the template analysis script and their insightful suggestions while setting up the task, Jan Hoffmann and Tobias Mühmer from the electrical and mechanical workshops of the Blumenbach Institute for Zoology and Anthropology of Göttingen University for building the behavioural arena, Parisa Jafari for conducting a subset the experiments, Prof. Dr. Andreas Stumpner for providing the chamber and advice for housing crickets and Christina Ahlbrecht for excellent animal care.

## AUTHOR CONTRIBUTIONS

Study design by S.B., S.L., and O.M.S. Prey capture data collection and experimental supervision by S.B. Formal analyses by S.B. and L.J.F.W.V. VWT with subregion-selective knock-down data collection by J.H. and X.H. Data interpretation by S.B., L.J.F.W.V., X.H., C.S., O.M.S. and S.L. Funding acquisition by O.M.S. and S.L. Original manuscript draft by S.B. and S.L. Reviewing and editing of manuscript by S.B., L.J.F.W.V., C.S., S.L. and O.M.S.

## DECLARATION OF INTERESTS

The authors declare no competing interests.

## DECLARATION OF GENERATIVE AI AND AI-ASSISTED TECHNOLOGIES

During the preparation of this work, the authors did not use any AI or AI-assisted tools.

## SUPPLEMENTAL INFORMATION

**Document S1. Figures S1–S3**

**Figure S1. WT mice are better hunters than KO mice even when inexperienced**

**Figure S2. Baseline immobility duration subtracted prey capture and detection times on test days**

**Figure S3. Predatory behaviour of WTs/KOs under darkness**

**Table S1. Prey capture test days (light trials) - Linear mixed effects analysis outputs**

**Table S2. Prey capture test days (dark trials) - Linear mixed effects analysis outputs**

**Video S1. Representative trial of a PSD-95 WT mouse with binocular vision**

**Video S2. Representative trial of a PSD-95 WT mouse with monocular vision**

**Video S3. Representative trial of a PSD-95 KO mouse with binocular vision**

**Video S4. Representative trial of a PSD-95 KO mouse with monocular vision**

## METHODS

### KEY RESOURCES TABLE

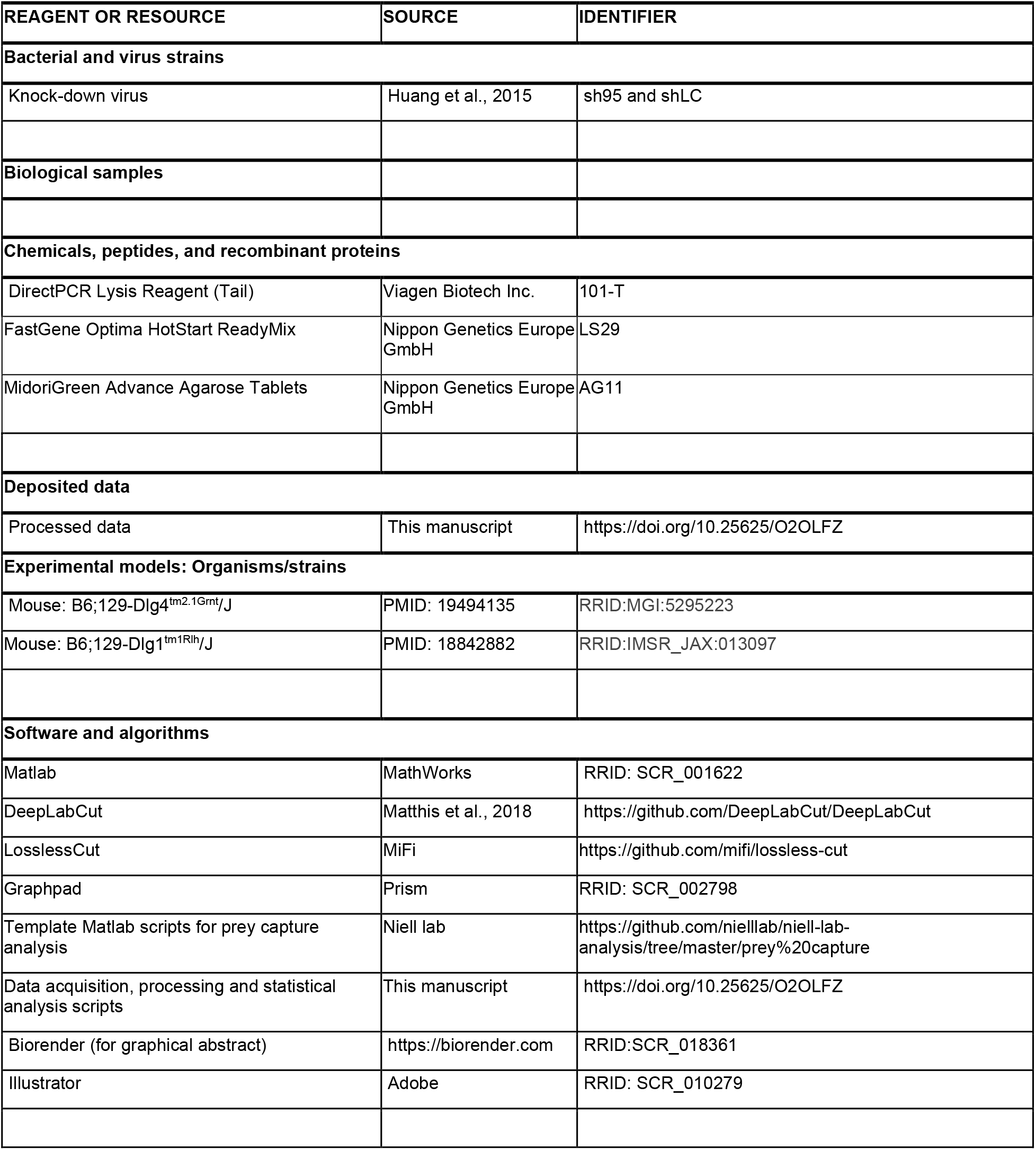

### EXPERIMENTAL MODEL AND STUDY PARTICIPANT DETAILS

All experimental procedures were approved by the local government: Niedersächsisches Landesamt für Verbraucherschutz und Lebensmittelsicherheit and Institutional Animal Care and Use Committee (IACUC) of the University of Pittsburgh. To generate Dlg4 +/+ and Dlg4 −/− transgenic mice used for the experiments, heterozygous female and male mice with a mixed 129SV/C57BL/6J background^38,110,111^, were bred at the central animal facility of the University Medical Center, Göttingen, Germany. To induce PSD-95 knock-down mice, C57BL/6J wild-type mice were obtained from the mouse colony of the central animal facility of the University of Pittsburgh, Pittsburgh, US. All mice were housed in standard cages (26×20×14 cm) with a 12/12 h light/dark cycle (light on at 08:00), with food pellets and water available ad libitum. For the visual water task experiment three cohorts of mice of both sexes were used all aged above P39, with MD performed prior to CP at p17-20: Cohort 1 consisted of 8 PSD-95/PSD-93 WT binocular, 6 PSD-95 WT monocular, 11 PSD-95 KO mice binocular and 4 PSD-95 KO mice with MD. Cohort 2 consisted of 12 C57BL/6J WT, 12 VC sh95 KD mice and 12 SC sh95 KD mice. Cohort 3 consisted of 6 C57BL/6J WT and 6 VC shLC control mice. All VWT experiments were conducted during the light cycle.

For the prey capture experiment 8 (3/5 ♀/♂) PSD-95 WT and 7 (1/6 ♀/♂) PSD-95 KO mice, all aged above P60 (mean±SD, WT: P220±85; KO: P217±70) were used. 9/10 female KO mice tested either did not catch crickets or failed to reach 100% capture success during the training phase and hence were not considered for further analyses. Medium-sized (2-3 cm) crickets (*Acheta domesticus*) of both sexes were sourced from Bugs-International GmbH and were maintained in a 12-hr reversed day-night cycle (light on at 22:00), such that the crickets’ night cycle coincided with the mice’s light cycle, which was when the experiments were conducted.

### METHOD DETAILS

#### Genotyping

Mouse tissue obtained from ear punctures were used for genotyping to perform a Polymerase chain reaction (PCR). Genomic DNA was extracted using DirectPCR Lysis Reagent (Viagen, 101-T) following the manufacturer’s instructions. PCR amplification was performed using FastGene Optima HotStart ReadyMix (Nippon Genetic Europe, LS29) using manufacturer’s instructions with the primer pair CAGGTGCTGCTGGAAGAAGG and CTACCCTGTGATCCAGAGCTG to detect both the WT and KO allele with sizes of 255 and 355 bp, respectively. PCR products were separated by electrophoresis on a 2% Midori^green^ advance agarose gel (Nippon genetic Europe, AG11) and visualized under cyan light.

#### Prey capture task

The general prey capture experimental and analysis protocol was adapted from previously published literature^13,14,19^ (Figure 1A-B). All trials in the behavioural arena (45 x 35 x 30 cm) were recorded using an overhead camera (60 fps; BFS-U3-32S4M). An array of alternating normal and near-infrared (NIR) LEDs were fixed on the lid of the arena to provide illumination under both light and dark conditions. The light trials were recorded under constant illumination (∼80 lx) along with infrared light to provide better image contrast. Trials under darkness were recorded using the NIR LEDs (∼880 nm) only. Prior to the start of the acclimatisation phase, mice were separated and singly housed throughout the experiment. The prey capture task consisted of a brief acclimatisation phase (days A1-A3) of 5-10min habituation with the experimenter and introduction of 1-2 crickets, kept overnight in the home cage from A1 onwards. At the end of A3, mice were introduced to the behavioural arena and allowed to habituate for 10min, after which the mice were put back to their home cages and food-restricted for ∼16h/day throughout the experimental timeline. During the training phase (after A3, starting with day D1), each mouse underwent 3-5 trials/day with one live cricket per trial. Each trial lasted for a maximum of 10 min and the arena was cleaned thoroughly with 70% ethanol in between each trial to eliminate any visual and/or olfactory cues from preceding trials. The training phase continued until an animal reached consistent hunting performance. The training phase continued until an animal reached consistent hunting performance. The maximum number of possible training days was limited to 15 days. Due to individual differences in the task, the total number of training days required to proceed to the test phase varied by subject. Due to high day-to-day variability, especially in KO hunting performance, hunting consistency was characterised by 100% capture success over at least the last two consecutive training days and median capture time differences not being greater than 15s between any two of the last four training days. The final test phase consisted of 2 days where the mice were first tested binocularly and then underwent monocular eye closure at the end of the 1st test day (D_n_) to be tested monocularly on the 2nd test day (D_n+1_), <24 h later. Both test days consisted of trials under both light and dark conditions, divided equally over 6-10 trails, in a pseudorandom order.

#### Monocular eye closure

After the binocular test day trials (D_n_), we closed the right eyes of the mice as described previously^32,34,38^. Mice were anaesthetised with 2% isoflurane in a mixture of O_2_:N_2_O (0.75:0.25), injected with carprofen (s.c., 5mg/kg), both eyelid margins were trimmed after which an antibiotic gel (Gentamicin-POS, Ursapharm) was applied. The eyelids were stitched shut using 2 mattress sutures (Ethicon, 7-0) and the mice were kept under a heating lamp for recovery (<5min). The monocularly deprived eye was inspected carefully to ensure it was closed before the monocular trials on the next day.

#### Knock-down of PSD-95

To induce a local KD of PSD-95 in V1 or SC, C57BL6/J mice were injected with either an AAV, expressing sh95 (sh95) or shLuciferase (shLC), as a negative control^32^, at P0. Adeno-associated viruses (AAV2/8) were produced as described previously in HEK293T cells with iodixanol gradient purification^32^. P0-P1 mouse pups were anaesthetized on ice for 5 min and immobilized with a holder on the injection table, where the surface was kept at 4 °C. The intersection of the sagittal and the lambdoid suture (lambda) was visually identified through the skin. Injections of AAV (13.4 nL each, 23 nL/s) were delivered at two positions bilaterally into V1 (from lambda, in mm: AP +0.1, ML ±1.25, DV −0.8 and AP +0.3, ML ±1.8, DV −0.8) or SC (from lambda, in mm: AP −0.2, ML ±0.3, DV −0.4), using a glass capillary with a Nanoject II microinjector (Drummond). Viral expression was confirmed in the extracted brain with a blue light torch and an orange filter.

#### Orientation discrimination

A modified version of the visual water task^39,41^ was used to quantify orientation discrimination as previously described^34^. Briefly, the apparatus was composed of a trapezoidal [118 (length) x 40 (height) x 80 (width at wide end) x 25 (width at narrow end) cm] water-filled chamber with the broader end containing 2 monitors (35 x 26 cm) placed side by side on the wall and partitioned by a barrier. Mice were trained to discriminate between a static vertical and horizontal grating of low spatial frequency (0.086 cycles/degree) presented on the two screens in random order. The vertical orientation was consistently rewarded by associating it with a submerged platform which the mouse learned to reach to escape from the water while the orthogonal (non-rewarded) stimulus was presented on the other monitor. After successful training (for the WT vs. KO mice experiment they had to reach 70% accuracy at the lowest possible orientation difference, and for the WT vs. KD experiment they had to reach ≥90% accuracy on at least two consecutive training sessions and complete ≥3 training sessions), the orientation difference was gradually reduced at 5° steps until performance dropped below 70%. Then the orientation difference was adjusted back by 15 ° and the mice were test again with reducing orientation difference. This procedure was repeated 3 times. The orientation discrimination threshold was considered the average of the three smallest orientation differences the animal successfully discriminated with at least 70% accuracy. For monocular groups, the right eye was sutured shut before the critical period (∼P17-20). The published binocular data points from Favaro et al. (2018) were included in the PSD-95 KO binocular group. The WT binocular group data consists of WTs from both PSD-95 and PSD-93 mouse lines.

#### Prey capture data acquisition and analysis

All videos were acquired using a custom Python workflow. Videos, where mice caught the cricket before the behaviour could be recorded, were discarded from subsequent analyses. The acquired videos used for data analyses were first manually trimmed (LosslessCut; https://github.com/mifi/lossless-cut) by marking the start and end of individual trials defined as the moment when the arena lid was closed and the end of individual trials was defined as the first instance when the mouse successfully caught the cricket. The trimmed videos were then corrected for lens distortion using custom software (OpenCV, Python). DeepLabCut^112,113^ was used to track different points of interest (ears, mouse head centre, mouse body, cricket) by training a ResNet-101 architecture and refining it 3 times by relabelling incorrectly tracked points. The resulting network yielded a test error of only 1.97 pixels (∼0.1 cm). Tracking quality was further refined by linearly interpolating over coordinates which had low DLC confidence (DLC likelihood <0.8) and/or when the Euclidean distance between adjacent x- and y-coordinates was higher than 5.5 cm (‘jump’ frames), occurring twice within 700ms. The head position vector of the mouse was calculated by taking the arctangent of the point in between the two ears (mouse head coordinate) and the target (cricket) vector was defined as the vector from the mouse head to the tracked cricket coordinate. The two vectors were then used to estimate the azimuth between the mouse and the cricket. Mouse speed and immobility were calculated using the defined marker for the mouse body. Immobility during baseline was defined as those phases during non-approach epochs where the mouse speed was <0.5cm/s. Arrest-like states during active approaches were defined as phases where the mouse speed was <2cm/s but >0.5cm/s. Mouse speeds during non-approach phases, when the mouse was not stationary (mouse speed>0.2cm/s), were considered as the baseline speed. Approach phases were defined as epochs where i) mouse speed was >6 cm/s, and ii) mouse-cricket distance (range) decreased by a rate of 7 cm/s and iii) the azimuth was <150 degrees^14,15,33^. Contacts were present when i) mouse-cricket distance was <3 cm and ii) azimuth was <90°. A successful contact given an approach occurred when the mouse-cricket distance was <3 cm within 250 ms after the termination of an approach epoch. The probability of a capture given a contact was defined as 1/total number of contacts in each trial.

### QUANTIFICATION AND STATISTICAL ANALYSIS

All statistical analyses, unless otherwise specified, were conducted on the pooled trials of each experimental group as previously described^14^. The assumptions, such as normal distribution were checked for each parameter. For most of the prey capture parameters, a linear mixed-effects model (LME) with restricted maximum likelihood (REML) was used to maintain consistency of the analyses. The model equation for the test days was defined as:

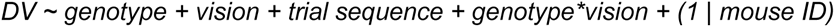

For the contact and capture time-dependent analysis, the model equation was similar, with the only difference being the addition of the underlined timestamp as dependent variable and random effect to the contact data:

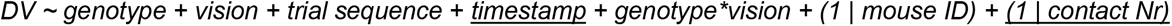

For the first training day analysis, the model equation was as follows:

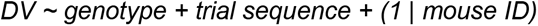

where, DV is the dependent variable, genotype (WT / KO) and vision (binocular/monocular) and trial sequence and timestamp in 50ms bins, are the fixed effect terms, the genotype*eye condition is the interaction term and (1 | mouse ID) or (1 | mouse ID) + (1 | contact Nr) are the random effect terms. The significance of the fixed effects and their interactions were assessed by Type II Wald χ^2^ tests and the post-hoc comparisons between the groups were computed using estimated marginal means for all simple main effect comparisons. Only relevant comparisons were considered and combined in one family for hypothesis testing. *p*-values were adjusted using the false discovery rate (FDR) method. The degrees of freedom for the comparisons were estimated using the Kenward-Roger method.

Orientation discrimination threshold differences between groups of Cohort 1 were assessed using an ordinary least squares (OLS) multiple linear regression model with the equation:

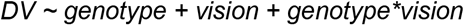

Cohorts 2 and 3 were combined and statistically assessed utilising an OLS multiple linear regression model with the equation:

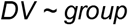

and estimated marginal means were used for multiple comparisons with the p-values corrected using the FDR method. No outlier correction was performed prior to statistical analyses, unless otherwise mentioned. The mixed-effects model, multiple linear regression model and related analyses were computed in R (using the stats, lme4 and emmeans packages) and the rest in either MATLAB (MathWorks) or GraphPad Prism. Effects were considered to be significant if *p*<0.05. The outputs of the LME analyses for the light and dark test days are given in *Tables S1* and *S2* of the supplementary materials.

## Notes

https://doi.org/10.25625/O2OLFZ

### Competing Interest Statement

The authors have declared no competing interest.

### Summary of Updates

First, we provide additional head movement analyses, specifically head angle in relation to body (head-body alignment) and in relation to cricket (azimuth) for the time (2 seconds) around prey contact under both light and dark conditions. Second, we provide additional experimental data demonstrating that knock-down of PSD-95 restricted to the primary visual cortex (V1) but not in superior colliculus (SC) significantly decreases orientation discrimination analyzed with the visual water task (VWT). We believe this new evidence better delineates the potential neuroanatomical locus of the PSD-95-associated perceptual deficits. Furthermore, we address the potential impact of PSD-95 KO learning deficits on our results. Moreover, in addition to binocular integration, we discuss alternative mechanisms that may underlie the impaired predation of PSD-95 KO mice with binocular vision. Finally, we address all other points raised by the reviewers (reviewer's comments available through eLife) through clearer exposition and reorganization of the manuscript.

