## Supplemental Figures for "PSD-95 drives binocular vision maturation critical for predation"

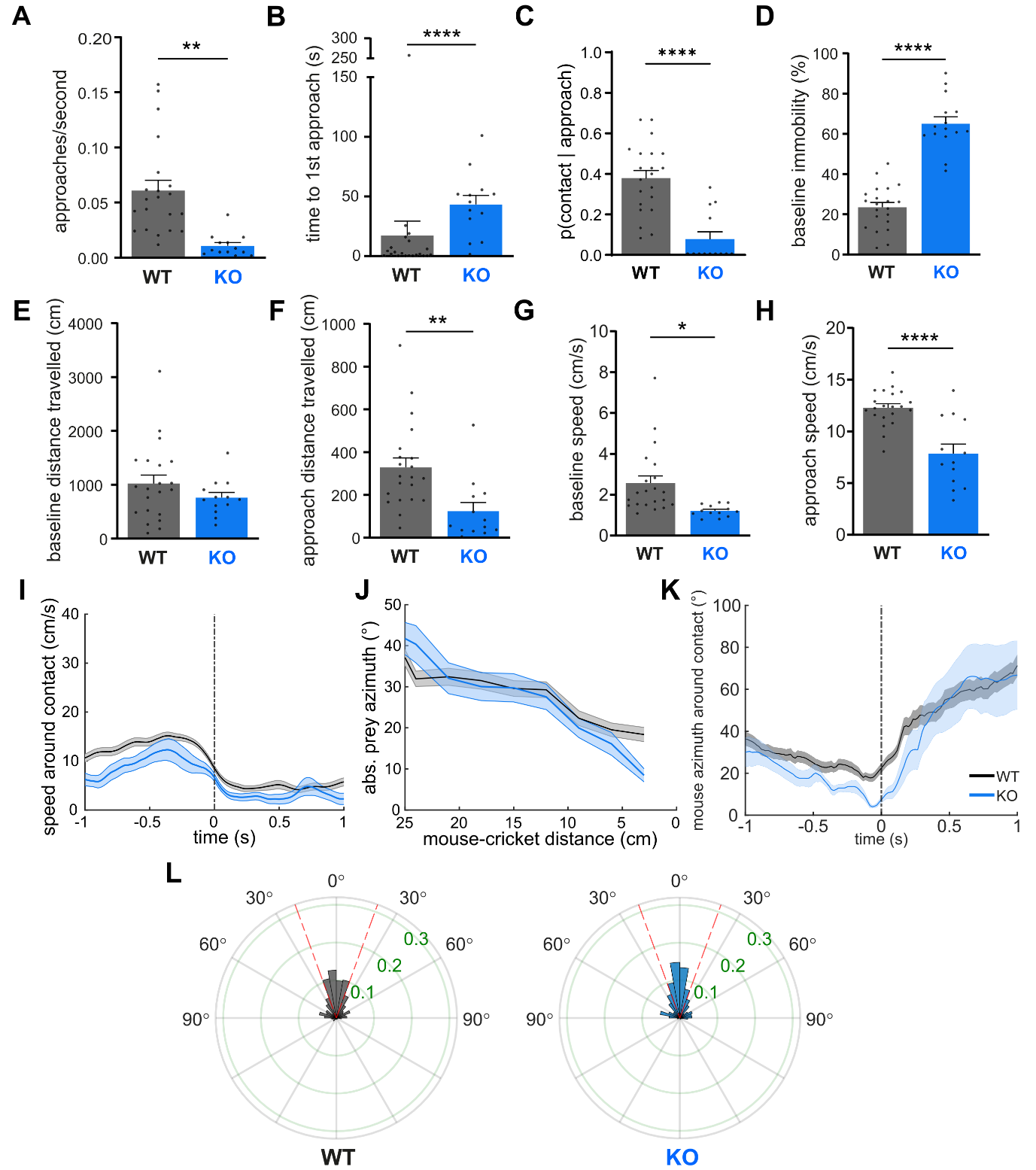

**Figure S1. Inexperienced WTs are better hunters than inexperienced KO mice. (A)** Prey approach frequency of PSD-95 WTs and KOs on the first day of cricket exposure in the arena. WT: 0.06±0.009, n=21; KO: 0.01±0.003, n=13. Linear mixed effects model, χ^2^_g_=9.157, p=0.002. **(B)** Time taken to initiate the first approach towards prey (immobility duration subtracted). WT: 17.16±12.11 s, n=21; KO: 43.28±7.562 s, n=13. Linear mixed effects model after outlier correction, χ^2^_g_=32.715, p<0.0001. **(C)** Probability to contact prey during an ongoing approach epoch. WT: 0.38±0.04, n=21; KO: 0.08±0.04, n=13. Linear mixed effects model, χ^2^_g_=19.536, p<0.0001. **(D)** Percentage of time mice stay immobile in a trial. WT: 23.48±2.38, n=21; KO: 65.06±3.43, n=15. Linear mixed effects model, χ^2^_g_=78.053, p<0.0001. **(E)** Distance covered by mice when not actively approaching prey. WT: 1024±156.2 cm, n=21; KO: 763.7±93.7 cm, n=13. Linear mixed effects model, χ^2^_g_=1.136, p=0.286. **(F)** Distance traversed during prey approach phases. WT: 328.6±44.05 cm, n=21; KO: 124.2±40.11 cm, n=13. Linear mixed effects model, χ^2^_g_=8.254, p=0.004. **(G)** Speed of mice when not stationary and not approaching prey. WT: 2.57±0.35, n=21; KO: 1.21±0.08, n=13. Linear mixed effects model, χ^2^_g_=4.424, p=0.035. **(H)** Speed of mice when actively approaching prey. WT: 12.29±0.37, n=21; KO: 7.84±0.92, n=13. Linear mixed effects model, χ^2^_g_=16.596, p<0.0001. **(I)** Speed of mice 1s before and after contact with prey. Dashed line indicates the time of contact. **(J)** Absolute mean prey azimuth as a function of mouse-cricket distance till contact during approach. **(K)** Absolute mean prey azimuth as a function of mouse-cricket distance from 1 second before and after contact. **(L)** Polar probability histograms (10º bins) of azimuth at the end of approach towards prey. Dashed red line indicates the binocular visual field (40º). Bars indicate mean±SEM of pooled trials from 7/5 WT/KO mice. Data points in **A-H** correspond to values in trials. Each data point in **D** corresponds to % of immobility in each trial respectively. Data points in **G** and **H** correspond to the median mouse speed in a trial. *p < 0.05, **p < 0.01, ****p < 0.0001.

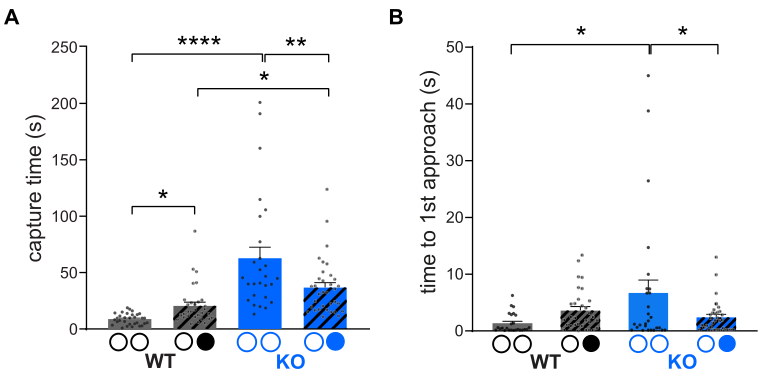
**Figure S2. Baseline immobility duration subtracted prey capture and detection times on test days. (A)** Prey capture time of WTs and KOs with binocular and monocular vision. WT binocular: 8.84±0.89 s, n=31; WT monocular: 20.76±3.04 s, n=32; KO binocular: 62.86±9.85 s, n=27; KO monocular: 36.96±4.24 s, n=34. **(B)** Latency to initiate first approach towards prey with and without binocular vision. WT binocular: 8.84±0.89 s, n=31; WT monocular: 3.63±0.66 s, n=31; KO binocular: 6.73±2.25 s, n=27; KO monocular: 2.42±0.49 s, n=34.

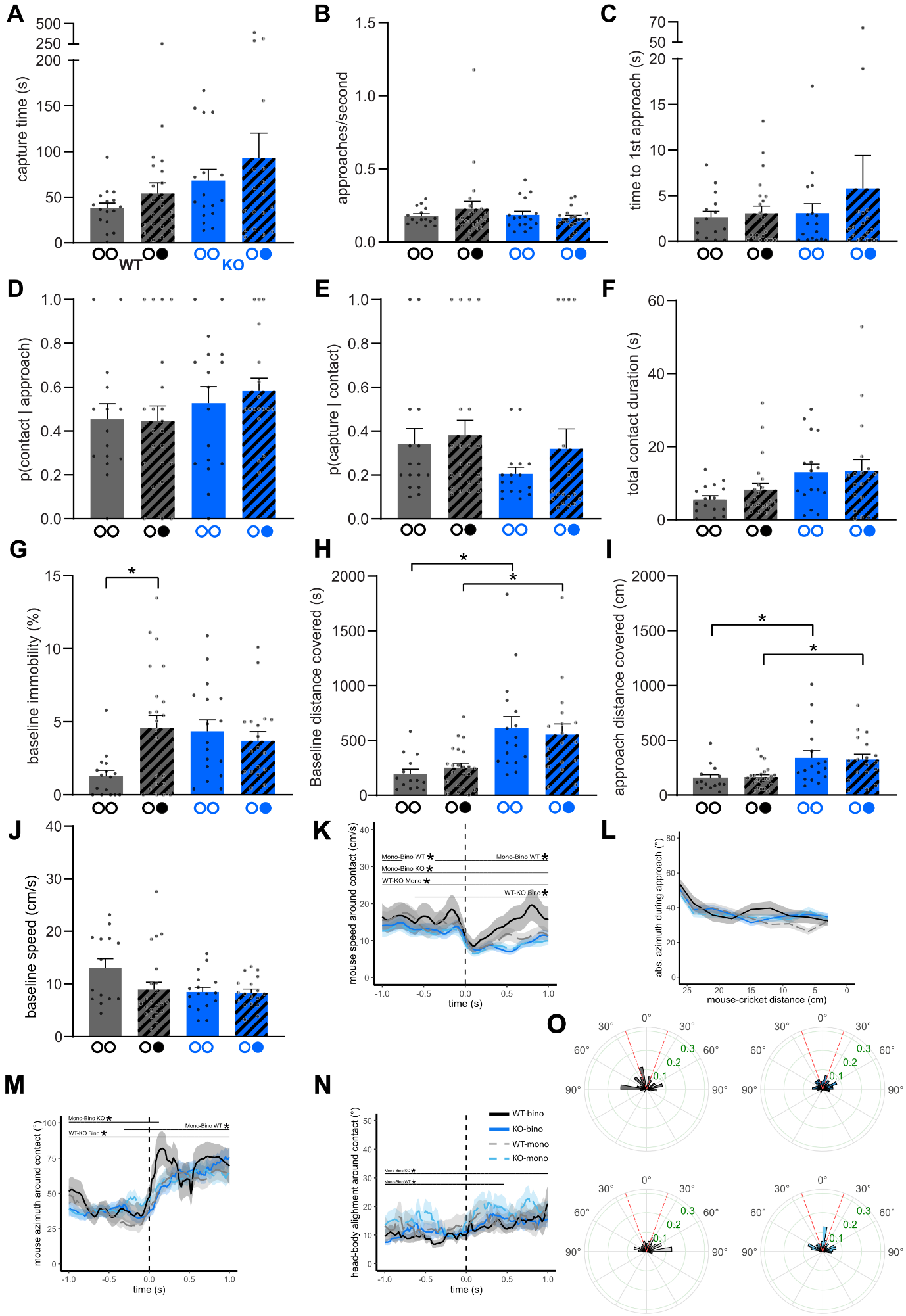

**Figure S3. Predatory behaviour of WTs/KOs under darkness. (A)** Time taken to capture prey. WT binocular: 37.72±5.57 s, n=16; WT monocular: 54.04±11.57 s, n=22; KO binocular: 68.25±12.25 s, n=17; KO monocular: 92.9±27.1 s, n=18. **(B)** Rate of approaches towards prey. WT binocular: 0.18±0.02, n=15; WT monocular: 0.23±0.05, n=22; KO binocular: 0.18±0.03, n=17; KO monocular: 0.17±0.02, n=18. **(C)** Latency to initiate the first approach sequence in a trial. WT binocular: 2.63±0.65 s, n=15; WT monocular: 3.06±0.77 s, n=22; KO binocular: 3.07±1.04 s, n=17; KO monocular: 5.79±3.59 s, n=18. **(D)** Probability of a contact given a successful approach. WT binocular: 0.45±0.07, n=15; WT monocular: 0.44±0.07, n=22; KO binocular: 0.53±0.07, n=17; KO monocular: 0.58±0.06, n=18. **(E)** Probability to capture prey. WT binocular: 0.34±0.07, n=16; WT monocular: 0.38±0.07, n=22; KO binocular: 0.21±0.03, n=17; KO monocular: 0.32±0.09, n=18. **(F)** Total time spent in contact with prey in a trial. WT binocular: 5.58±1 s, n=16; WT monocular: 8.28±1.66 s, n=22; KO binocular: 13.03±2.21 s, n=17; KO monocular: 13.4±3.08 s, n=18. **(G)** Baseline immobility of mice. WT binocular: 1.31±0.38%, n=16; WT monocular: 4.58±0.88%, n=22; KO binocular: 4.36±0.77%, n=17; KO monocular: 3.71±0.63%, n=18. **(H)** Distance travelled by mouse at baseline. WT binocular: 195.6±42.74 cm, n=14; WT monocular: 253.3±40.72 cm, n=22; KO binocular: 614.7±104.1 cm, n=17; KO monocular: 555.1±95.79 cm, n=18. **(I)** Distance travelled by mouse during approach phases. WT binocular: 154.1±31.23 cm, n=14; WT monocular: 165.1±21.53 cm, n=22; KO binocular: 341.4±64.31 cm, n=17; KO monocular: 325.7±48.85 cm, n=18. **(J)** Baseline speed of mice. WT binocular: 13.01±1.76 cm/s, n=14; WT monocular: 8.94±1.41 cm/s, n=22; KO binocular: 8.5±0.87 cm/s, n=17; KO monocular: 8.37±0.68 cm/s, n=18. **(K)** Mouse speed 1s before and after contact with prey. Lines and shaded areas indicate mean±SEM. Dashed line indicates the time of contact. **(L)** Prey azimuth as a function of mouse-cricket distance till the end of a successful approach. **(M)** Mean absolute azimuth 1s before and after contact with prey. Lines and shaded areas indicate mean±SEM. Dashed line indicates the time of a contact. **(N)** Head-to-body alignment of mice around ±1 s of contact with prey, with body and head completely aligned at an angle of 0. Lines and shaded areas indicate mean±SEM. Dashed line indicates the time of a contact. **(O)** Polar histogram of azimuth at the end of successful approaches. Dashed red line indicates the binocular visual field (±20º). Bars indicate mean±SEM of pooled trials from 4/6 WT and 5/4 KO mice under binocular and monocular conditions respectively. Each data point in **G** corresponds to % of immobility in each trial. Data points in **H** correspond to the median mouse speed in a trial. *p < 0.05.

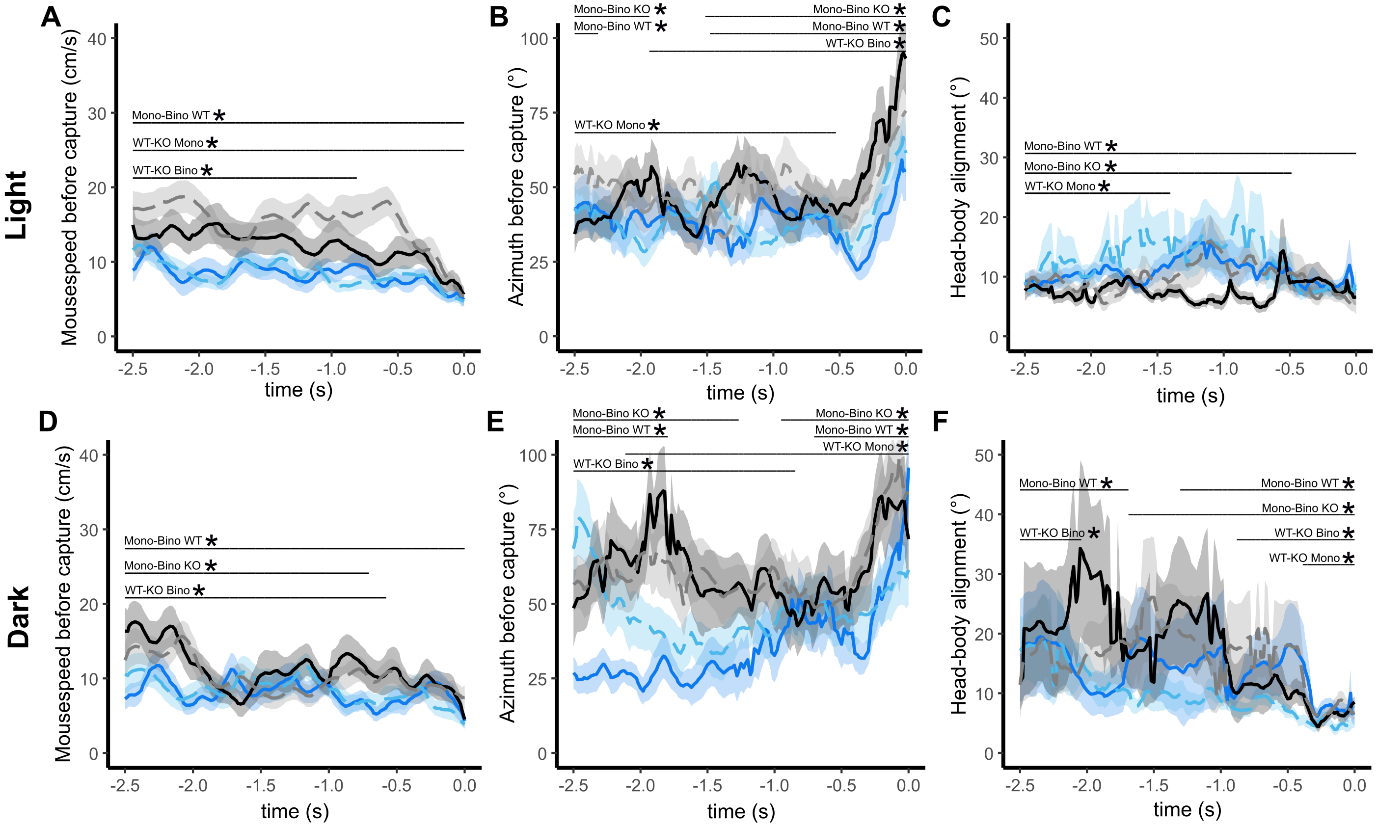

**Figure S4. Predatory behaviour of WTs/KOs before capture.**

**(A)** Mouse speed 2.5s before capturing the prey in light trials. **(B)** Mean absolute azimuth 2.5s before and after contact with prey in light trials. **(C)** Head-to-body alignment of mice around 2.5s before capturing the prey in light trials, with body and head completely aligned at an angle of 0. **(D)** Mouse speed 2.5s before capturing the prey in dark trials. **(E)** Mean absolute azimuth 2.5s before and after contact with prey in dark trials. **(F)** Head-to-body alignment of mice around 2.5s before capturing the prey in dark trials, with body and head completely aligned at an angle of 0. Lines and shaded areas indicate mean±SEM.
